# Diagnostic prediction tools for bacteraemia caused by 3rd generation cephalosporin-resistant Enterobacteriaceae in suspected bacterial infections: a nested case-control study

**DOI:** 10.1101/120550

**Authors:** Wouter C. Rottier, Cornelis H. van Werkhoven, Yara R.P. Bamberg, J. Wendelien Dorigo-Zetsma, Ewoudt M. van de Garde, Babette C. van Hees, Jan A.J.W. Kluytmans, Emile M. Kuck, Paul D. van der Linden, Jan M. Prins, Steven F.T. Thijsen, Annelies Verbon, Bart J.M. Vlaminckx, Heidi S.M. Ammerlaan, Marc J.M. Bonten

## Abstract

**Objectives:** Current guidelines for empirical antibiotic treatment poorly predict the presence of 3rd generation cephalosporin resistant Enterobacteriaceae (3GC-R EB) as a cause of infection, thereby increasing unnecessary carbapenem use. We aimed to develop diagnostic scoring systems to better predict the presence of 3GC-R EB as a cause of bacteraemia.

**Methods:** A retrospective nested case-control study was performed that included patients ≥18 years in whom blood cultures were obtained and intravenous antibiotics were initiated. Each patient with 3GC-R EB bacteraemia was matched to four control infection episodes within the same hospital, based on blood culture date and onset location (community or hospital). Starting from 32 described clinical risk factors at infection onset, selection strategies were used to derive scoring systems for the probability of community- and hospital-onset 3GC-R EB bacteraemia.

**Results:** 3GC-R EB bacteraemia occurred in 90 of 22,506 (0.4%) community-onset and in 82 of 8,110 (1.0%) hospital-onset infections, and these cases were matched to 360 community-onset and 328 hospital-onset control episodes, respectively. The derived community-onset and hospital-onset scoring system consisted of 6 and 9 predictors, respectively, with c-statistics of 0.807 (95% confidence interval 0.756-0.855) and 0.842 (0.794-0.887). With selected score cutoffs, the models identified 3GC-R EB bacteraemia with equal sensitivity as existing guidelines, but reduced the proportion of patients classified as at risk for 3GC-R EB bacteraemia (i.e. eligible for empiric carbapenem therapy) with 40% in patients with community-onset and 49% in patients with hospital-onset infection.

**Conclusions:** These prediction rules for 3GC-R EB bacteraemia may reduce unnecessary empiric carbapenem use.

## Introduction

As a consequence of the emergence of infections caused 3^rd^ generation cephalosporin (3GC) resistant Enterobacteriaceae (3GC-R EB; in this manuscript used synonymously with extended-spectrum β-lactamase (ESBL) producing Enterobacteriaceae), physicians are increasingly faced with the question which patients need empiric antibiotic treatment covering these pathogens. Hence, patients and physicians might benefit from prediction rules for 3GC-R EB. Although risk factors for carriage of ESBL-producing Enterobacteriaceae at hospital admission [1–4], and factors distinguishing ESBL- and carbapenemase-producing Enterobacteriaceae as a cause of bacteraemia have been determined [5–8], there are no prediction rules for identifying 3GC-R EB as a cause of bacteraemia at the time that empiric therapy must be started.

Current Dutch empiric treatment guidelines designate patients at risk of infection caused by 3GC-R EB based on prior colonization or infection with 3GC-R EB or on prior exposure to cephalosporins or fluoroquinolones, as these were identified as risk factors in patients with bacteraemia caused by these pathogens [9]. As carbapenems are the treatment of choice for 3GC-R EB, adherence to these guidelines may stimulate overuse of these antibiotics. Indeed, applying these recommendations for all patients needing empiric antibiotic treatment in a population with a pre-test probability for 3GC-R EB of 0.7%, revealed that 19% of all patients were classified as at risk for 3GC-R EB infection and thus eligible for empiric carbapenem therapy (referred to as test positivity rate), while at the same time only 50% of all patients with 3GC-R EB bacteraemia would be classified as at risk (referred to as sensitivity) [10]. Only using prior identification of 3GC-R EB carriage as risk factor, would reduce the test positivity rate to 4%, at the cost of a reduction in sensitivity to 42%.

We aimed to develop prediction rules to better identify, among patients needing intravenous empiric antibiotic therapy, those being infected with 3GC-R EB. We were specifically interested in the balance between sensitivity and test positivity rate. In this derivation study, we compared these quantities to those of the two basic strategies introduced above, which rely on prior identification alone (*prior identification model*), or in combination with prior exposure to cephalosporins and fluoroquinolones (*two-predictor model*). We focused on predicting 3GC-R EB bacteraemia, as these infections can be objectively assessed in retrospect, and an immediate start with appropriate antibiotics is indicated. We decided to derive separate prediction rules for community-onset and hospital-onset infections, as we assumed that factors driving spread of 3GC-R EB within these two settings are distinct.

## Methods

### Setting and patients

This was a retrospective nested case-control study involving 8 hospitals, of which 3 university hospitals, in the Netherlands. Between January 1^st^ 2008 and December 31^st^ 2010, we included all consecutive patients of 18 years of age or older in whom a blood culture was obtained and intravenous broad-spectrum β-lactam antibiotics (i.e. not penicillin or flucloxacillin), aminoglycosides, and/or fluoroquinolones were started on the day of the blood culture or the day after, irrespective of duration. Patients receiving any of the eligible antibiotics on the day of blood culture obtainment were excluded if these had been initiated prior to this day (see Supplementary Table 1 for illustrating examples). In addition, patients with 3GC-R EB bacteraemia in the year prior were excluded, as it was assumed that treating physicians would always provide therapy aimed at these organisms in case of renewed infection. Patients could be included more than once, if a subsequent episode complied with in- and exclusion criteria. Additional information on hospital characteristics, study periods, and databases used in each of the hospitals is provided in Supplementary Table 2.

Infection episodes were separated into two cohorts: the community-onset cohort comprised episodes in which the first blood culture was collected during the first three calendar days of hospitalization, and the hospital-onset cohort consisted of episodes in which blood cultures were obtained later during hospitalization.

The causative pathogen of each episode was based on the results of blood cultures obtained on the day that antibiotics were started and the day before. In both cohorts, the case population comprised all consecutive infection episodes with 3GC-R EB bacteraemia (see Supplementary Table 2 for definition of 3GC resistance in each of the hospitals). We estimated that a study period of three years in the participating hospitals would yield 100 patients with 3GC-R EB bacteraemia in both cohorts, which would allow initial logistic regression with 10 variables, based on the 10 events per variable recommendation [11].

The control population was defined as all other infection episodes, including non-bacteraemic episodes and episodes with blood cultures yielding non-resistant Enterobacteriaceae, other bacteria or fungi. From this population four controls were matched to each case, a ratio chosen because of minimal gains in statistical power with more controls [12]. Controls were matched on hospital, being in the community or hospital-onset cohort, and being closest in time to the blood culture day of the case episode.

Due to its retrospective nature, the Dutch Medical Research Involving Human Subjects Act did not apply to this study. In each of the participating hospitals, applicable local guidelines for noninterventional studies were followed. In accordance with Dutch regulations, informed consent was waived for the study. Reporting of this study was in accordance with the TRIPOD Statement [13,14].

### Data collection

All selected cases and controls were subjected to chart review to obtain information that was considered available at the moment that the initial antibiotics were prescribed (referred to as infection onset). Blinding for the outcome during chart review was not considered feasible. Please refer to Supplementary Table 3 for an overview of all collected variables.

### Statistical analysis

Two separate prediction models were constructed, one for community-onset and one for hospital-onset infections. Data analyses were performed in R (version 3.2.2) [15], including packages *mice* 2.25 [16], *rms* 4.5-0 [17], *pROC* 1.8 [18], and *xtable* 1.8-2 [19]. Descriptive analyses of predictors were based on non-missing data only. Some variables were aggregated because of high correlation, low prevalence, and/or similar associations with the outcome (indicated in Table 1). Additionally, the number of categories for suspected sources was reduced to four by combining categories with low frequencies into a single remaining group (original categories in Supplementary Table 3), and categories for antibiotic use were created based on prevalence and assumed predictive power for 3GC-R EB infection. Twenty imputed datasets were created to deal with missing values during the modelling stage. In the Supplementary Material, missing data patterns and the exact imputation procedure are described.

**Table 1.**
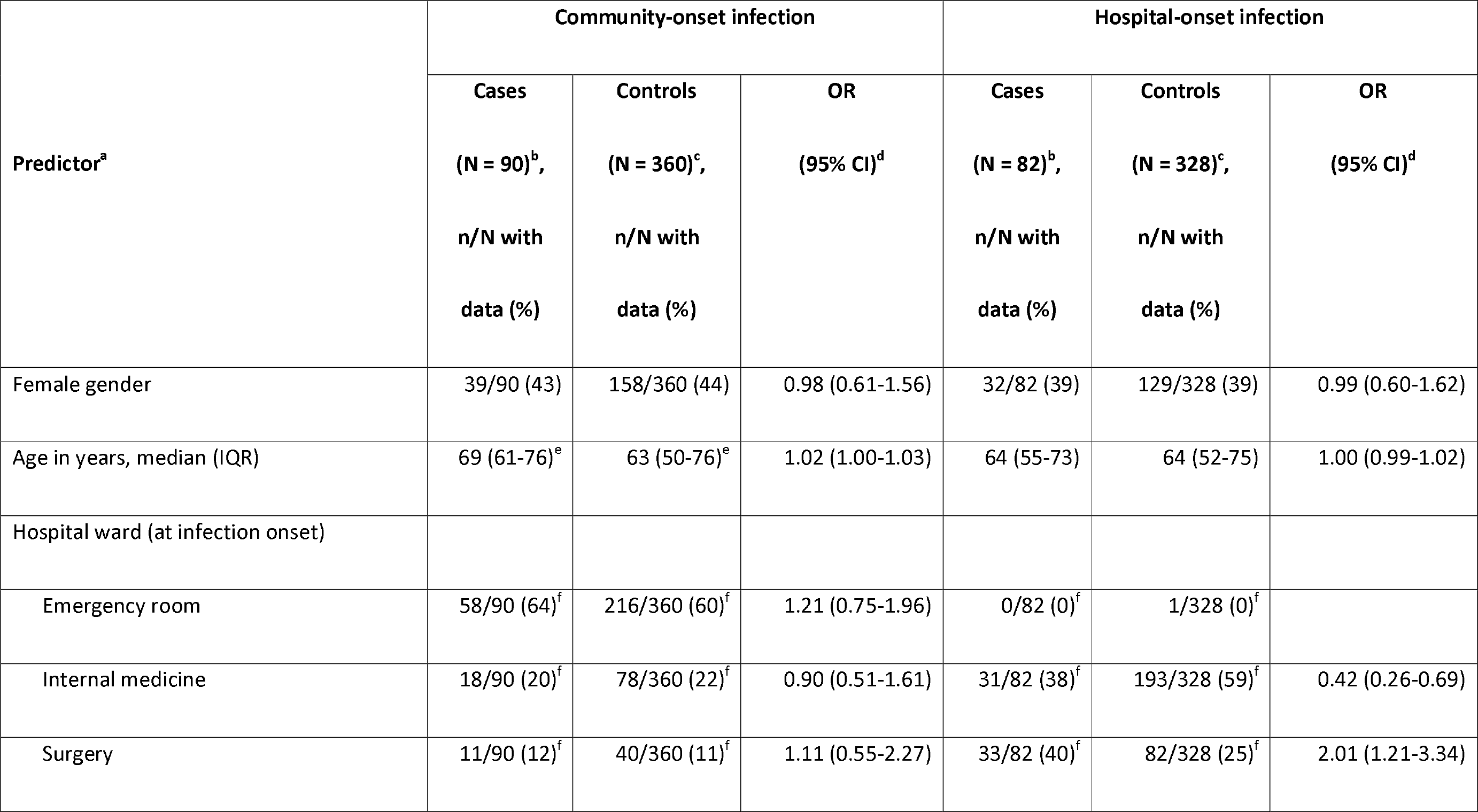
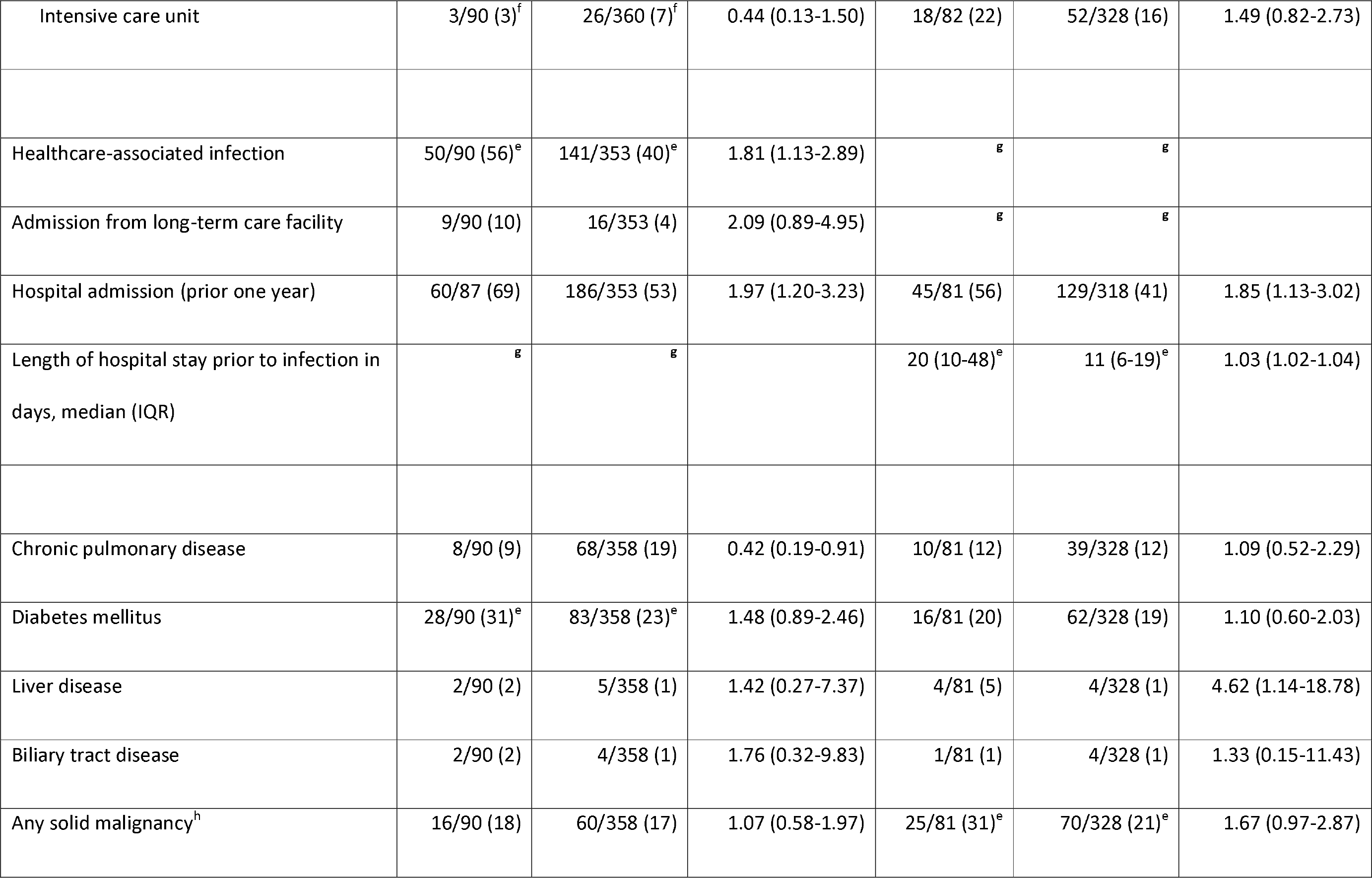
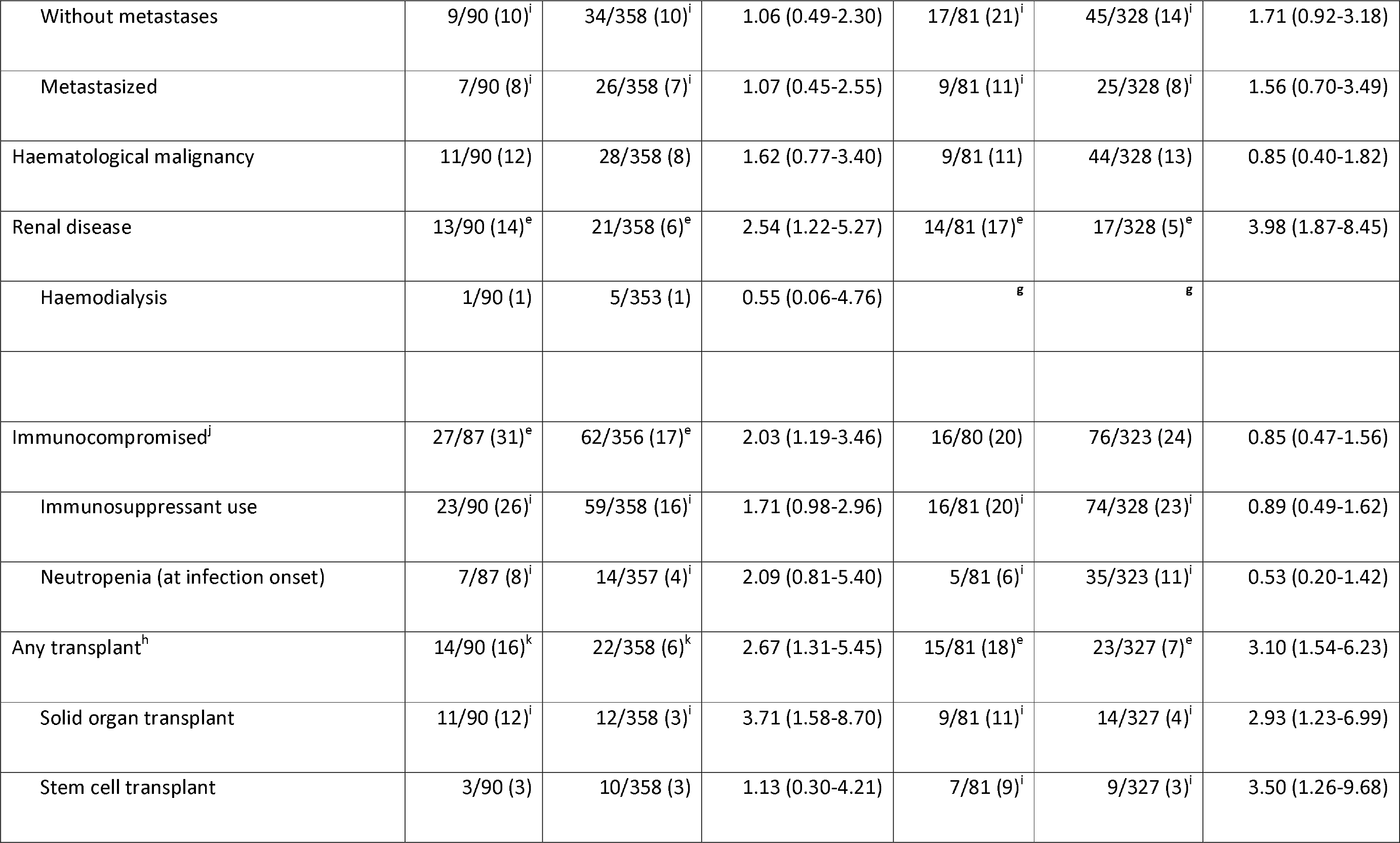
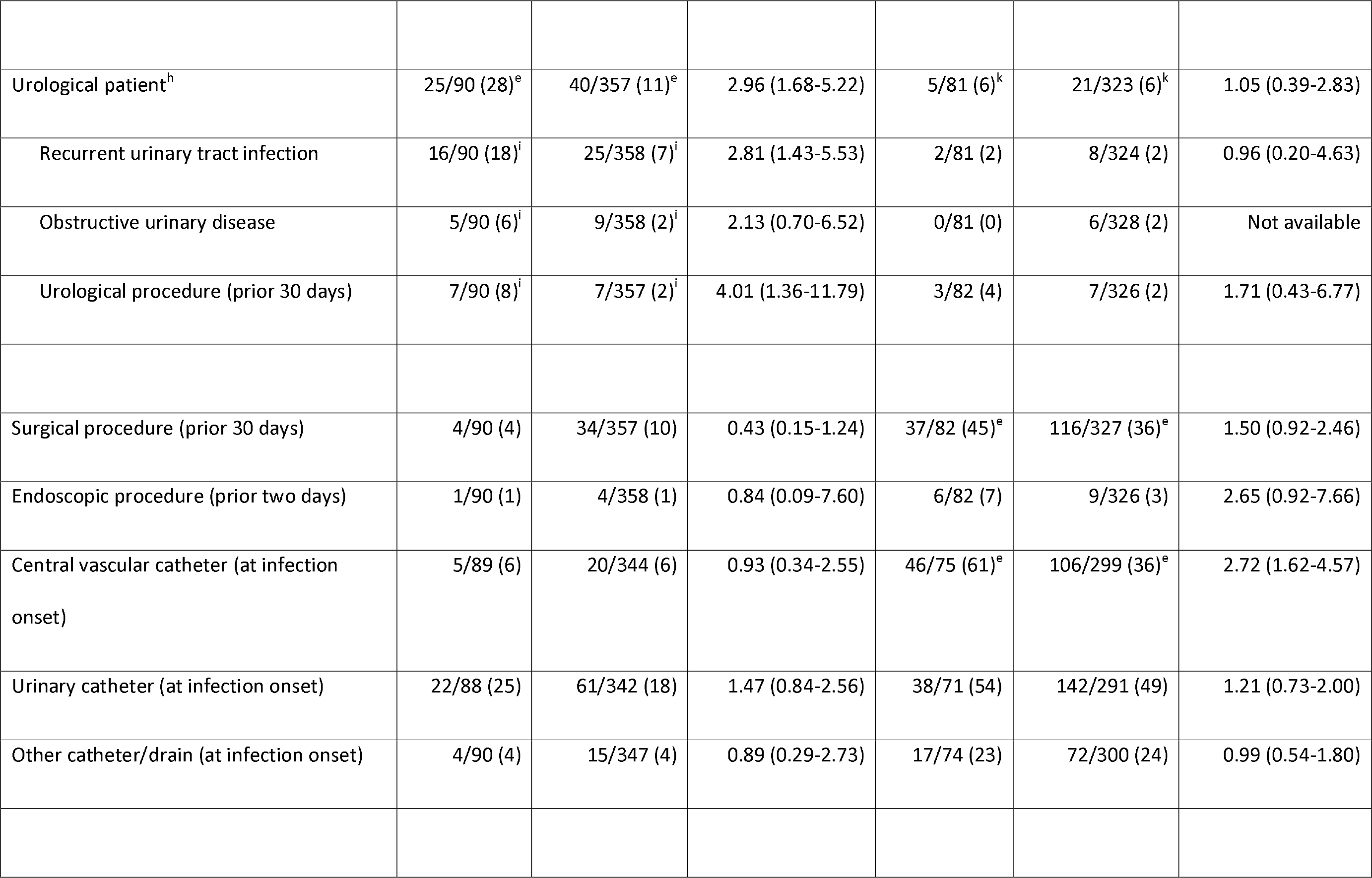
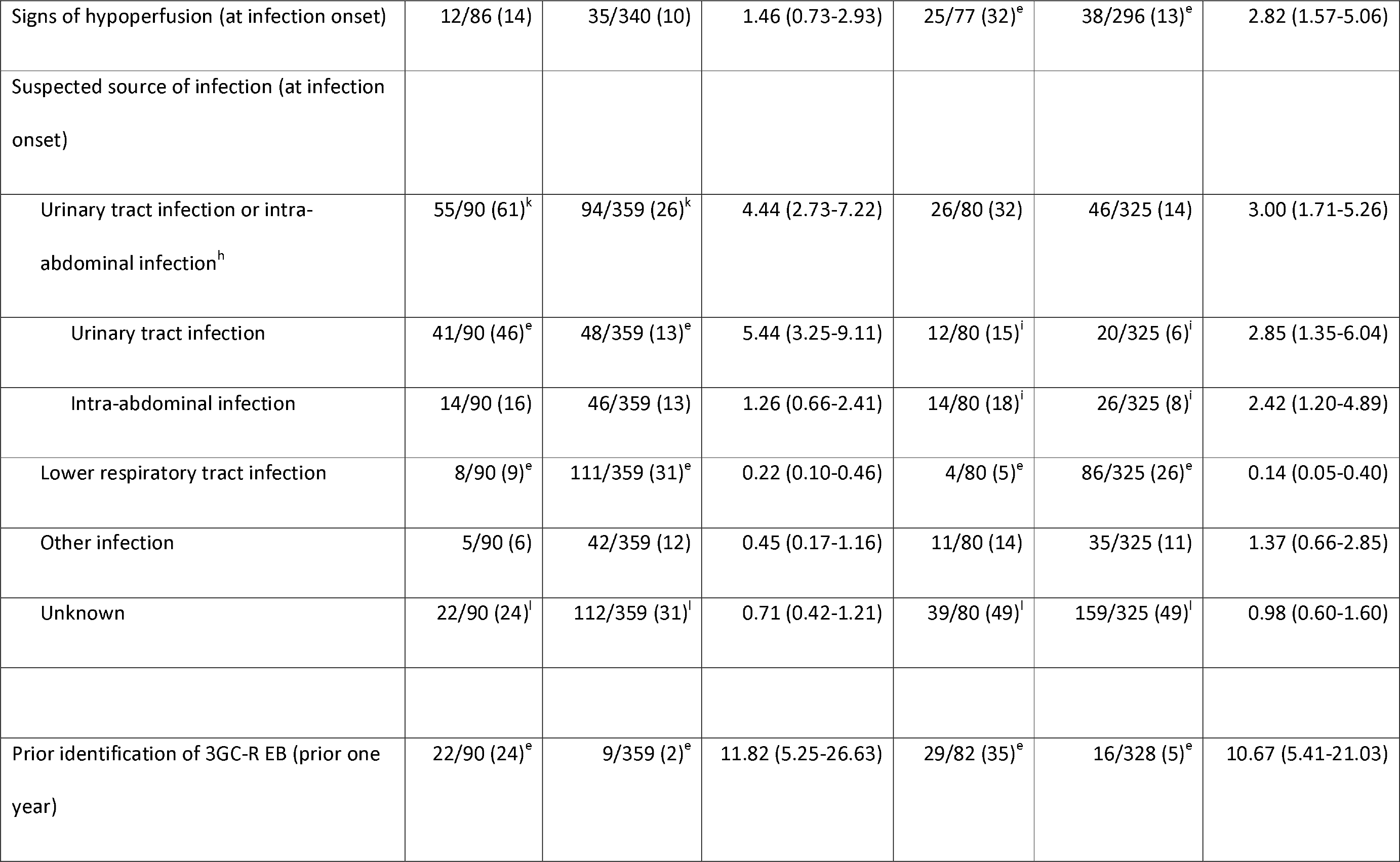
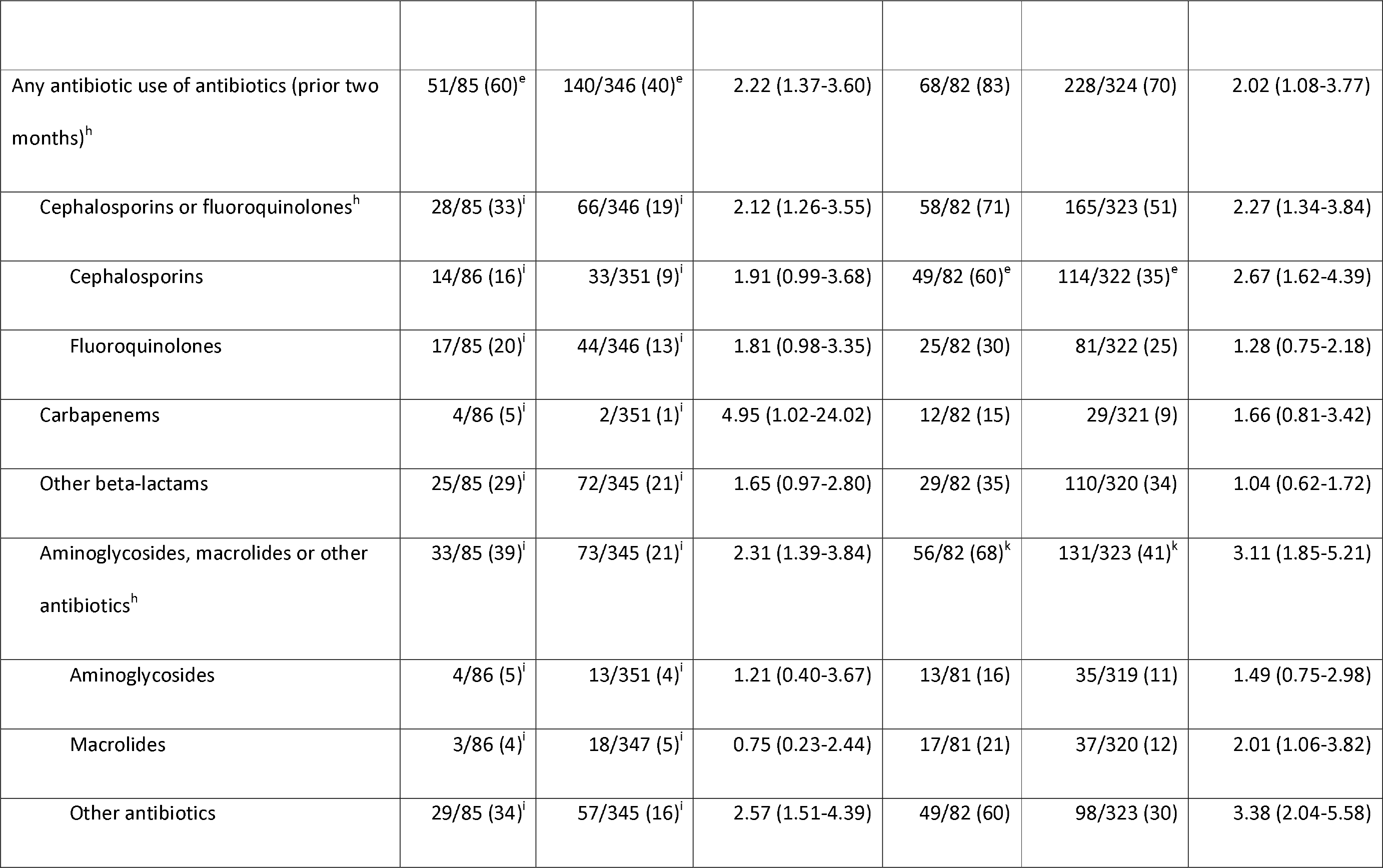
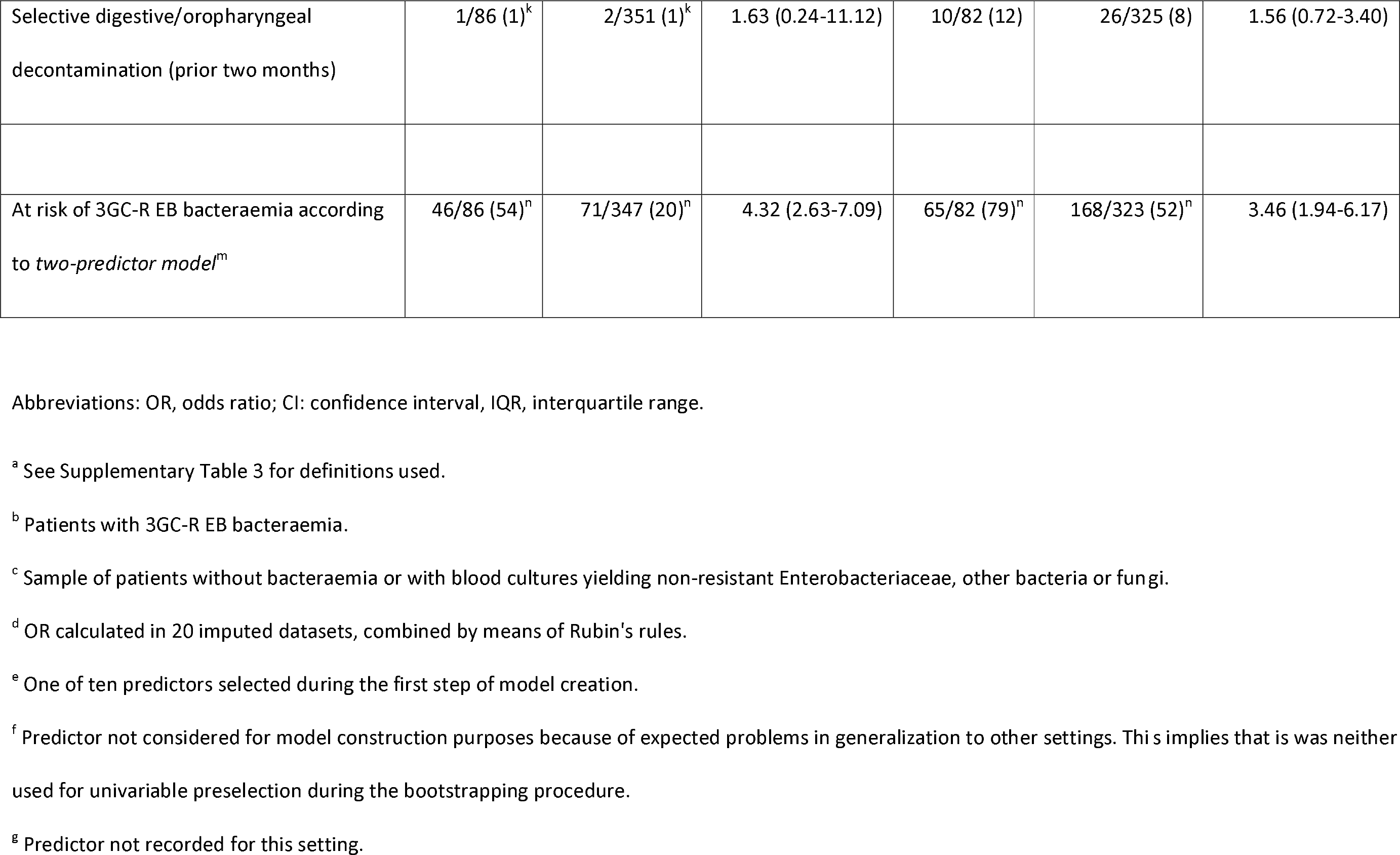
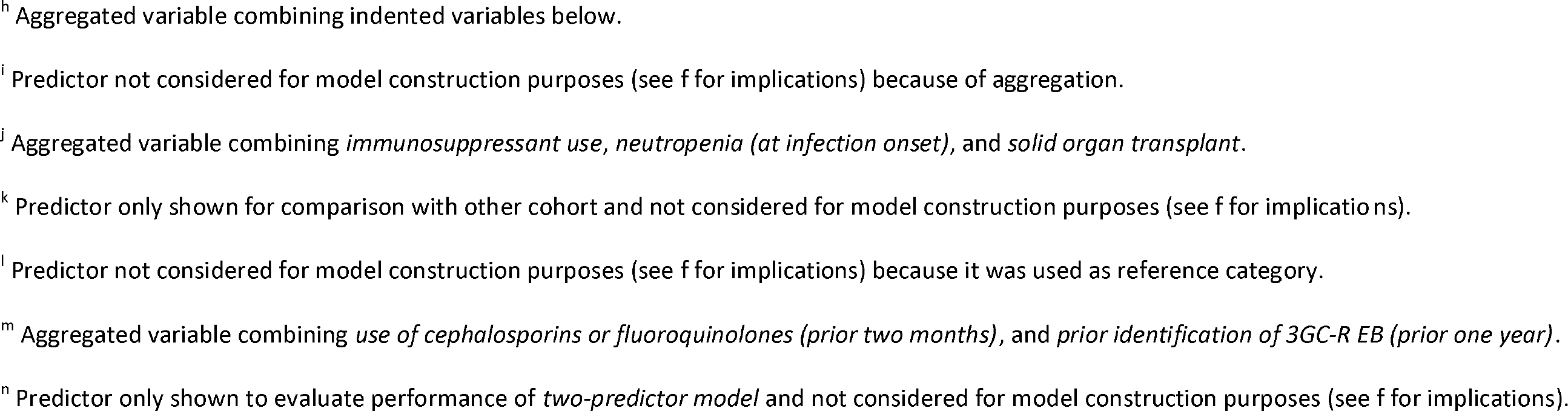
Clinical characteristics of cases and controls from both the community-onset and hospital-onset cohort.

| **Predictor<sup>a</sup>** | **Community-onset infection** |  |  | **Hospital-onset infection** |  |  |
| --- | --- | --- | --- | --- | --- | --- |
|  | **Cases**<br>**(N = 90)<sup>b</sup>,**<br>**n/N with**<br>**data (%)** | **Controls**<br>**(N = 360)<sup>c</sup>,**<br>**n/N with**<br>**data (%)** | **OR**<br>**(95% CI)<sup>d</sup>** | **Cases**<br>**(N = 82)<sup>b</sup>,**<br>**n/N with**<br>**data (%)** | **Controls**<br>**(N = 328)<sup>c</sup>,**<br>**n/N with**<br>**data (%)** | **OR**<br>**(95% CI)<sup>d</sup>** |
| Female gender | 39/90 (43) | 158/360 (44) | 0.98 (0.61-1.56) | 32/82 (39) | 129/328 (39) | 0.99 (0.60-1.62) |
| Age in years, median (IQR) | 69 (61-76)<sup>e</sup> | 63 (50-76)<sup>e</sup> | 1.02 (1.00-1.03) | 64 (55-73) | 64 (52-75) | 1.00 (0.99-1.02) |
| Hospital ward (at infection onset) |  |  |  |  |  |  |
| Emergency room | 58/90 (64)<sup>f</sup> | 216/360 (60)<sup>f</sup> | 1.21 (0.75-1.96) | 0/82 (0)<sup>f</sup> | 1/328 (0)<sup>f</sup> |  |
| Internal medicine | 18/90 (20)<sup>f</sup> | 78/360 (22)<sup>f</sup> | 0.90 (0.51-1.61) | 31/82 (38)<sup>f</sup> | 193/328 (59)<sup>f</sup> | 0.42 (0.26-0.69) |
| Surgery | 11/90 (12)<sup>f</sup> | 40/360 (11)<sup>f</sup> | 1.11 (0.55-2.27) | 33/82 (40)<sup>f</sup> | 82/328 (25)<sup>f</sup> | 2.01 (1.21-3.34) |
| Intensive care unit | 3/90 (3)<sup>f</sup> | 26/360 (7)<sup>f</sup> | 0.44 (0.13-1.50) | 18/82 (22) | 52/328 (16) | 1.49 (0.82-2.73) |
| Healthcare-associated infection | 50/90 (56)<sup>e</sup> | 141/353 (40)<sup>e</sup> | 1.81 (1.13-2.89) | <sup>g</sup> | <sup>g</sup> |  |
| Admission from long-term care facility | 9/90 (10) | 16/353 (4) | 2.09 (0.89-4.95) | <sup>g</sup> | <sup>g</sup> |  |
| Hospital admission (prior one year) | 60/87 (69) | 186/353 (53) | 1.97 (1.20-3.23) | 45/81 (56) | 129/318 (41) | 1.85 (1.13-3.02) |
| Length of hospital stay prior to infection in days, median (IQR) | <sup>g</sup> | <sup>g</sup> |  | 20 (10-48)<sup>e</sup> | 11 (6-19)<sup>e</sup> | 1.03 (1.02-1.04) |
| Chronic pulmonary disease | 8/90 (9) | 68/358 (19) | 0.42 (0.19-0.91) | 10/81 (12) | 39/328 (12) | 1.09 (0.52-2.29) |
| Diabetes mellitus | 28/90 (31)<sup>e</sup> | 83/358 (23)<sup>e</sup> | 1.48 (0.89-2.46) | 16/81 (20) | 62/328 (19) | 1.10 (0.60-2.03) |
| Liver disease | 2/90 (2) | 5/358 (1) | 1.42 (0.27-7.37) | 4/81 (5) | 4/328 (1) | 4.62 (1.14-18.78) |
| Biliary tract disease | 2/90 (2) | 4/358 (1) | 1.76 (0.32-9.83) | 1/81 (1) | 4/328 (1) | 1.33 (0.15-11.43) |
| Any solid malignancy<sup>h</sup> | 16/90 (18) | 60/358 (17) | 1.07 (0.58-1.97) | 25/81 (31)<sup>e</sup> | 70/328 (21)<sup>e</sup> | 1.67 (0.97-2.87) |
| Without metastases | 9/90 (10)<sup>i</sup> | 34/358 (10)<sup>i</sup> | 1.06 (0.49-2.30) | 17/81 (21)<sup>i</sup> | 45/328 (14)<sup>i</sup> | 1.71 (0.92-3.18) |
| Metastasized | 7/90 (8)<sup>i</sup> | 26/358 (7)<sup>i</sup> | 1.07 (0.45-2.55) | 9/81 (11)<sup>i</sup> | 25/328 (8)<sup>i</sup> | 1.56 (0.70-3.49) |
| Haematological malignancy | 11/90 (12) | 28/358 (8) | 1.62 (0.77-3.40) | 9/81 (11) | 44/328 (13) | 0.85 (0.40-1.82) |
| Renal disease | 13/90 (14)<sup>e</sup> | 21/358 (6)<sup>e</sup> | 2.54 (1.22-5.27) | 14/81 (17)<sup>e</sup> | 17/328 (5)<sup>e</sup> | 3.98 (1.87-8.45) |
| Haemodialysis | 1/90 (1) | 5/353 (1) | 0.55 (0.06-4.76) | <sup>g</sup> | <sup>g</sup> |  |
| Immunocompromised<sup>j</sup> | 27/87 (31)<sup>e</sup> | 62/356 (17)<sup>e</sup> | 2.03 (1.19-3.46) | 16/80 (20) | 76/323 (24) | 0.85 (0.47-1.56) |
| Immunosuppressant use | 23/90 (26)<sup>i</sup> | 59/358 (16)<sup>i</sup> | 1.71 (0.98-2.96) | 16/81 (20)<sup>i</sup> | 74/328 (23)<sup>i</sup> | 0.89 (0.49-1.62) |
| Neutropenia (at infection onset) | 7/87 (8)<sup>i</sup> | 14/357 (4)<sup>i</sup> | 2.09 (0.81-5.40) | 5/81 (6)<sup>i</sup> | 35/323 (11)<sup>i</sup> | 0.53 (0.20-1.42) |
| Any transplant<sup>h</sup> | 14/90 (16)<sup>k</sup> | 22/358 (6)<sup>k</sup> | 2.67 (1.31-5.45) | 15/81 (18)<sup>e</sup> | 23/327 (7)<sup>e</sup> | 3.10 (1.54-6.23) |
| Solid organ transplant | 11/90 (12)<sup>i</sup> | 12/358 (3)<sup>i</sup> | 3.71 (1.58-8.70) | 9/81 (11)<sup>i</sup> | 14/327 (4)<sup>i</sup> | 2.93 (1.23-6.99) |
| Stem cell transplant | 3/90 (3) | 10/358 (3) | 1.13 (0.30-4.21) | 7/81 (9)<sup>i</sup> | 9/327 (3)<sup>i</sup> | 3.50 (1.26-9.68) |
| Urological patient<sup>h</sup> | 25/90 (28)<sup>e</sup> | 40/357 (11)<sup>e</sup> | 2.96 (1.68-5.22) | 5/81 (6)<sup>k</sup> | 21/323 (6)<sup>k</sup> | 1.05 (0.39-2.83) |
| Recurrent urinary tract infection | 16/90 (18)<sup>i</sup> | 25/358 (7)<sup>i</sup> | 2.81 (1.43-5.53) | 2/81 (2) | 8/324 (2) | 0.96 (0.20-4.63) |
| Obstructive urinary disease | 5/90 (6)<sup>i</sup> | 9/358 (2)<sup>i</sup> | 2.13 (0.70-6.52) | 0/81 (0) | 6/328 (2) | Not available |
| Urological procedure (prior 30 days) | 7/90 (8)<sup>i</sup> | 7/357 (2)<sup>i</sup> | 4.01 (1.36-11.79) | 3/82 (4) | 7/326 (2) | 1.71 (0.43-6.77) |
| Surgical procedure (prior 30 days) | 4/90 (4) | 34/357 (10) | 0.43 (0.15-1.24) | 37/82 (45)<sup>e</sup> | 116/327 (36)<sup>e</sup> | 1.50 (0.92-2.46) |
| Endoscopic procedure (prior two days) | 1/90 (1) | 4/358 (1) | 0.84 (0.09-7.60) | 6/82 (7) | 9/326 (3) | 2.65 (0.92-7.66) |
| Central vascular catheter (at infection onset) | 5/89 (6) | 20/344 (6) | 0.93 (0.34-2.55) | 46/75 (61)<sup>e</sup> | 106/299 (36)<sup>e</sup> | 2.72 (1.62-4.57) |
| Urinary catheter (at infection onset) | 22/88 (25) | 61/342 (18) | 1.47 (0.84-2.56) | 38/71 (54) | 142/291 (49) | 1.21 (0.73-2.00) |
| Other catheter/drain (at infection onset) | 4/90 (4) | 15/347 (4) | 0.89 (0.29-2.73) | 17/74 (23) | 72/300 (24) | 0.99 (0.54-1.80) |
| Signs of hypoperfusion (at infection onset) | 12/86 (14) | 35/340 (10) | 1.46 (0.73-2.93) | 25/77 (32)<sup>e</sup> | 38/296 (13)<sup>e</sup> | 2.82 (1.57-5.06) |
| Suspected source of infection (at infection onset) |  |  |  |  |  |  |
| Urinary tract infection or intra-abdominal infection<sup>h</sup> | 55/90 (61)<sup>k</sup> | 94/359 (26)<sup>k</sup> | 4.44 (2.73-7.22) | 26/80 (32) | 46/325 (14) | 3.00 (1.71-5.26) |
| Urinary tract infection | 41/90 (46)<sup>e</sup> | 48/359 (13)<sup>e</sup> | 5.44 (3.25-9.11) | 12/80 (15)<sup>i</sup> | 20/325 (6)<sup>i</sup> | 2.85 (1.35-6.04) |
| Intra-abdominal infection | 14/90 (16) | 46/359 (13) | 1.26 (0.66-2.41) | 14/80 (18)<sup>i</sup> | 26/325 (8)<sup>i</sup> | 2.42 (1.20-4.89) |
| Lower respiratory tract infection | 8/90 (9)<sup>e</sup> | 111/359 (31)<sup>e</sup> | 0.22 (0.10-0.46) | 4/80 (5)<sup>e</sup> | 86/325 (26)<sup>e</sup> | 0.14 (0.05-0.40) |
| Other infection | 5/90 (6) | 42/359 (12) | 0.45 (0.17-1.16) | 11/80 (14) | 35/325 (11) | 1.37 (0.66-2.85) |
| Unknown | 22/90 (24)<sup>i</sup> | 112/359 (31)<sup>i</sup> | 0.71 (0.42-1.21) | 39/80 (49)<sup>i</sup> | 159/325 (49)<sup>i</sup> | 0.98 (0.60-1.60) |
| Prior identification of 3GC-R EB (prior one year) | 22/90 (24)<sup>e</sup> | 9/359 (2)<sup>e</sup> | 11.82 (5.25-26.63) | 29/82 (35)<sup>e</sup> | 16/328 (5)<sup>e</sup> | 10.67 (5.41-21.03) |
| Any antibiotic use of antibiotics (prior two months)<sup>h</sup> | 51/85 (60)<sup>e</sup> | 140/346 (40)<sup>e</sup> | 2.22 (1.37-3.60) | 68/82 (83) | 228/324 (70) | 2.02 (1.08-3.77) |
| Cephalosporins or fluoroquinolones<sup>h</sup> | 28/85 (33)<sup>i</sup> | 66/346 (19)<sup>i</sup> | 2.12 (1.26-3.55) | 58/82 (71) | 165/323 (51) | 2.27 (1.34-3.84) |
| Cephalosporins | 14/86 (16)<sup>i</sup> | 33/351 (9)<sup>i</sup> | 1.91 (0.99-3.68) | 49/82 (60)<sup>e</sup> | 114/322 (35)<sup>e</sup> | 2.67 (1.62-4.39) |
| Fluoroquinolones | 17/85 (20)<sup>i</sup> | 44/346 (13)<sup>i</sup> | 1.81 (0.98-3.35) | 25/82 (30) | 81/322 (25) | 1.28 (0.75-2.18) |
| Carbapenems | 4/86 (5)<sup>i</sup> | 2/351 (1)<sup>i</sup> | 4.95 (1.02-24.02) | 12/82 (15) | 29/321 (9) | 1.66 (0.81-3.42) |
| Other beta-lactams | 25/85 (29)<sup>i</sup> | 72/345 (21)<sup>i</sup> | 1.65 (0.97-2.80) | 29/82 (35) | 110/320 (34) | 1.04 (0.62-1.72) |
| Aminoglycosides, macrolides or other antibiotics<sup>h</sup> | 33/85 (39)<sup>i</sup> | 73/345 (21)<sup>i</sup> | 2.31 (1.39-3.84) | 56/82 (68)<sup>k</sup> | 131/323 (41)<sup>k</sup> | 3.11 (1.85-5.21) |
| Aminoglycosides | 4/86 (5)<sup>i</sup> | 13/351 (4)<sup>i</sup> | 1.21 (0.40-3.67) | 13/81 (16) | 35/319 (11) | 1.49 (0.75-2.98) |
| Macrolides | 3/86 (4)<sup>i</sup> | 18/347 (5)<sup>i</sup> | 0.75 (0.23-2.44) | 17/81 (21) | 37/320 (12) | 2.01 (1.06-3.82) |
| Other antibiotics | 29/85 (34)<sup>i</sup> | 57/345 (16)<sup>i</sup> | 2.57 (1.51-4.39) | 49/82 (60) | 98/323 (30) | 3.38 (2.04-5.58) |
| Selective digestive/oropharyngeal decontamination (prior two months) | 1/86 (1)<sup>k</sup> | 2/351 (1)<sup>k</sup> | 1.63 (0.24-11.12) | 10/82 (12) | 26/325 (8) | 1.56 (0.72-3.40) |
| At risk of 3GC-R EB bacteraemia according to *two-predictor model*<sup>m</sup> | 46/86 (54)<sup>n</sup> | 71/347 (20)<sup>n</sup> | 4.32 (2.63-7.09) | 65/82 (79)<sup>n</sup> | 168/323 (52)<sup>n</sup> | 3.46 (1.94-6.17) |

Abbreviations: OR, odds ratio; CI: confidence interval, IQR, interquartile range.
<sup>a</sup> See Supplementary Table 3 for definitions used.
<sup>b</sup> Patients with 3GC-R EB bacteraemia.
<sup>c</sup> Sample of patients without bacteraemia or with blood cultures yielding non-resistant Enterobacteriaceae, other bacteria or fungi.
<sup>d</sup> OR calculated in 20 imputed datasets, combined by means of Rubin's rules.
<sup>e</sup> One of ten predictors selected during the first step of model creation.
<sup>f</sup> Predictor not considered for model construction purposes because of expected problems in generalization to other settings. This implies that it was neither used for univariable preselection during the bootstrapping procedure.
<sup>g</sup> Predictor not recorded for this setting.
<sup>h</sup> Aggregated variable combining indented variables below.
<sup>i</sup> Predictor not considered for model construction purposes (see f for implications) because of aggregation.
<sup>j</sup> Aggregated variable combining *immunosuppressant use*, *neutropenia (at infection onset)*, and *solid organ transplant*.
<sup>k</sup> Predictor only shown for comparison with other cohort and not considered for model construction purposes (see f for implications).
<sup>l</sup> Predictor not considered for model construction purposes (see f for implications) because it was used as reference category.
<sup>m</sup> Aggregated variable combining *use of cephalosporins or fluoroquinolones (prior two months)*, and *prior identification of 3GC-R EB (prior one year)*.

Starting from 32 potential predictors, the first step of model creation involved selection of ten relevant predictors based on (1) observing the strength of their association with 3GC-R EB bacteraemia (without statistical hypothesis testing), and (2) considerations related to coverage of the entire spectrum of known risk factors for 3GC-R EB, and (3) ease-of-use of any resulting model. The second step involved removing redundant variables from the model, which was performed by backward stepwise logistic regression analysis until all remaining predictors had p-values < 0.2 in the Wald test (pooled from 20 imputed datasets by means of Rubin’s rules) [20]. Continuous predictors were initially introduced into models with restricted cubic spline functions with three knots to allow for non-linear associations. Finally, we evaluated by means of the Akaike’s Information Criterion (AIC) if simplification to a linear predictor was possible.

Regression coefficients of the final models were pooled over imputed datasets by means of Rubin’s rules and shrunk according to model optimism (see description further on). Furthermore, developing a model in a case-control study artificially increases the prevalence of the outcome, which means that predicted probabilities generated by the model do not reflect true probabilities within the full cohorts. Test positivity rates, and positive and negative predictive values are similarly affected. Therefore, intercepts of the models were adjusted for the sampling fraction of the controls, and controls were weighted by the inverse of the sampling fraction, as previously described [21]. All quantities presented in this paper reflect the values within the original full cohorts.

Calibration of the predicted and observed probabilities was visually inspected for separate imputed datasets. All other performance parameters were averaged over the imputed datasets. Discrimination was assessed with the area under the curve for receiver operating characteristic curves (referred to as C-statistic). Sensitivity, specificity and positive and negative predictive values, and test positivity rate (i.e. fraction of the population classified as at risk of 3GC-R EB bacteraemia) were calculated for different cutoffs of the predicted risk. These model performance characteristics were compared to those of the *prior identification model* and *two-predictor model*. In the *prior identification model*, patients with identification of 3GC-R EB in the year prior to the infection episode were classified as test-positive. In the *two-predictor model*, also patients with cephalosporin or fluoroquinolone use during the prior two months were considered test-positive.

A simplified score was created by multiplying the regression coefficients with a constant, followed by rounding to easy-to-use values. Performance of this score was determined similarly.

### Estimation of model optimism

Optimism results from the fact that models are developed on a population sample and suffer from overfitting, which jeopardizes generalizability to other populations, including future patients for which a model will be used [22]. By means of a bootstrapping technique, the expected performance loss (e.g. lower sensitivity, specificity, and predictive values, and altered test positivity rate) when applying the model within the total population is quantified. For the two regression models, optimism was estimated by creating 2000 bootstrap samples, creating a new prediction model for each of these samples, and comparing the model’s performance in the original and bootstrapped data. Optimism was estimated for model coefficients, derived odds ratios and C-statistics. During the same procedure, the expected overestimation of sensitivity and underestimation of test positivity rate due to optimism was quantified by applying a probability cutoff above which patients are classified as test-positive. For this evaluation, the probability cutoff was selected such that sensitivity corresponded to the *two-predictor model*. In the Supplementary Material, technical details of the bootstrapping procedure are presented.

## Results

Probabilities of 3GC-R EB bacteraemia were 0.4% (n = 90) for the community-onset infection cohort (22,506 episodes) and 1.0% (n = 82) for the hospital-onset infection cohort (8,110 episodes) (Figure 1). These case populations were matched to 360 community-onset control episodes and 328 hospital-onset control episodes (Table 1). Multiple selection of individual patients, albeit with different episodes, as case and/or control were allowed and occurred 8 times within the community-onset, and 9 times within the hospital-onset dataset. Isolated pathogens from blood cultures and initial antibiotic therapy are presented in Supplementary Tables 4 and 5.

**Figure 1.**
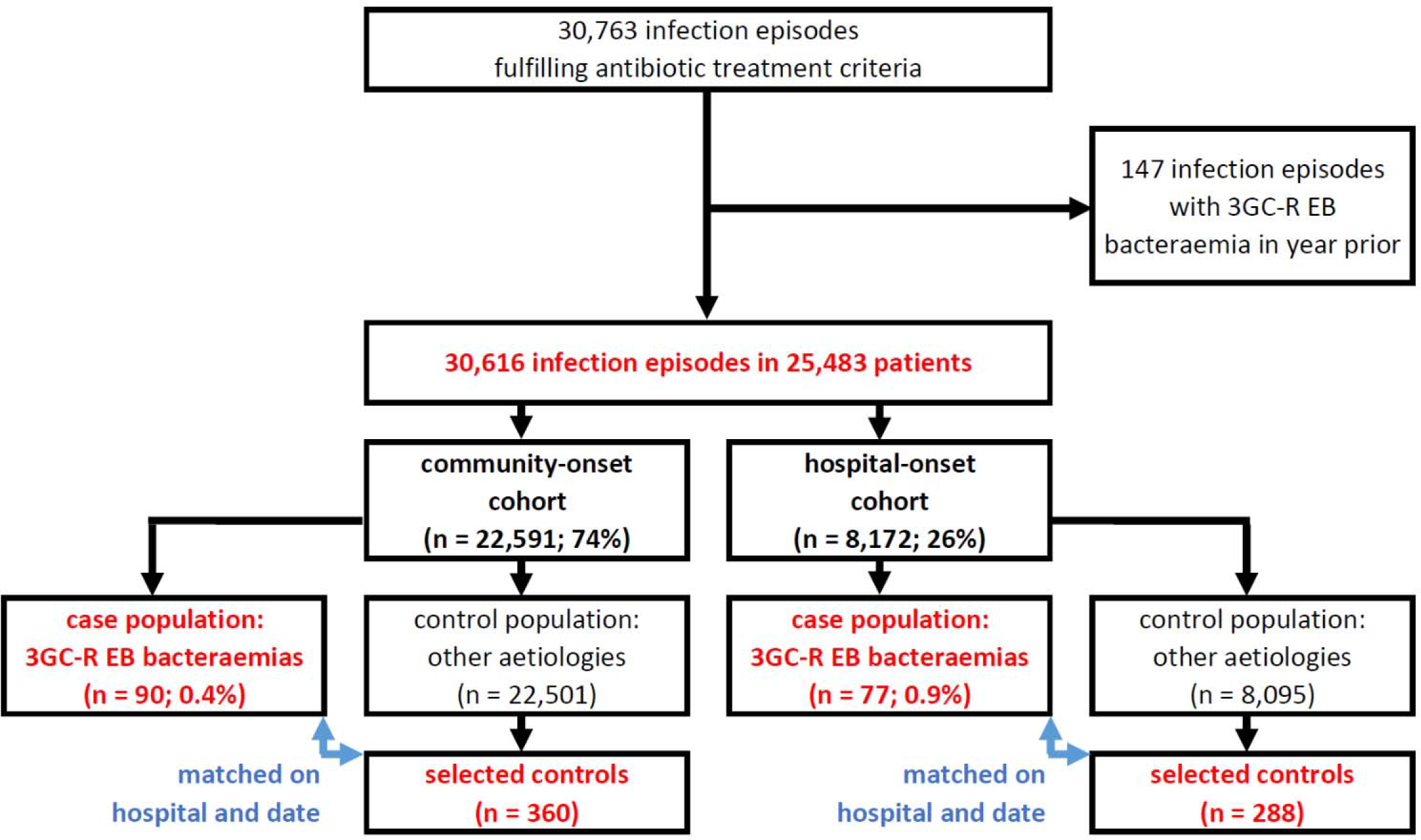
Patient flowchart.

### Community-onset infection

The prediction model for 3GC-R EB bacteraemia in community-onset infection consisted of six variables (Table 2). It showed adequate discrimination (C-statistic = 0.808 (95% CI 0.756-0.855), also after correction for optimism (C-statistic = 0.775 (95% CI 0.705-0.839)), and calibration (Supplementary Figure 1).

**Table 2.**
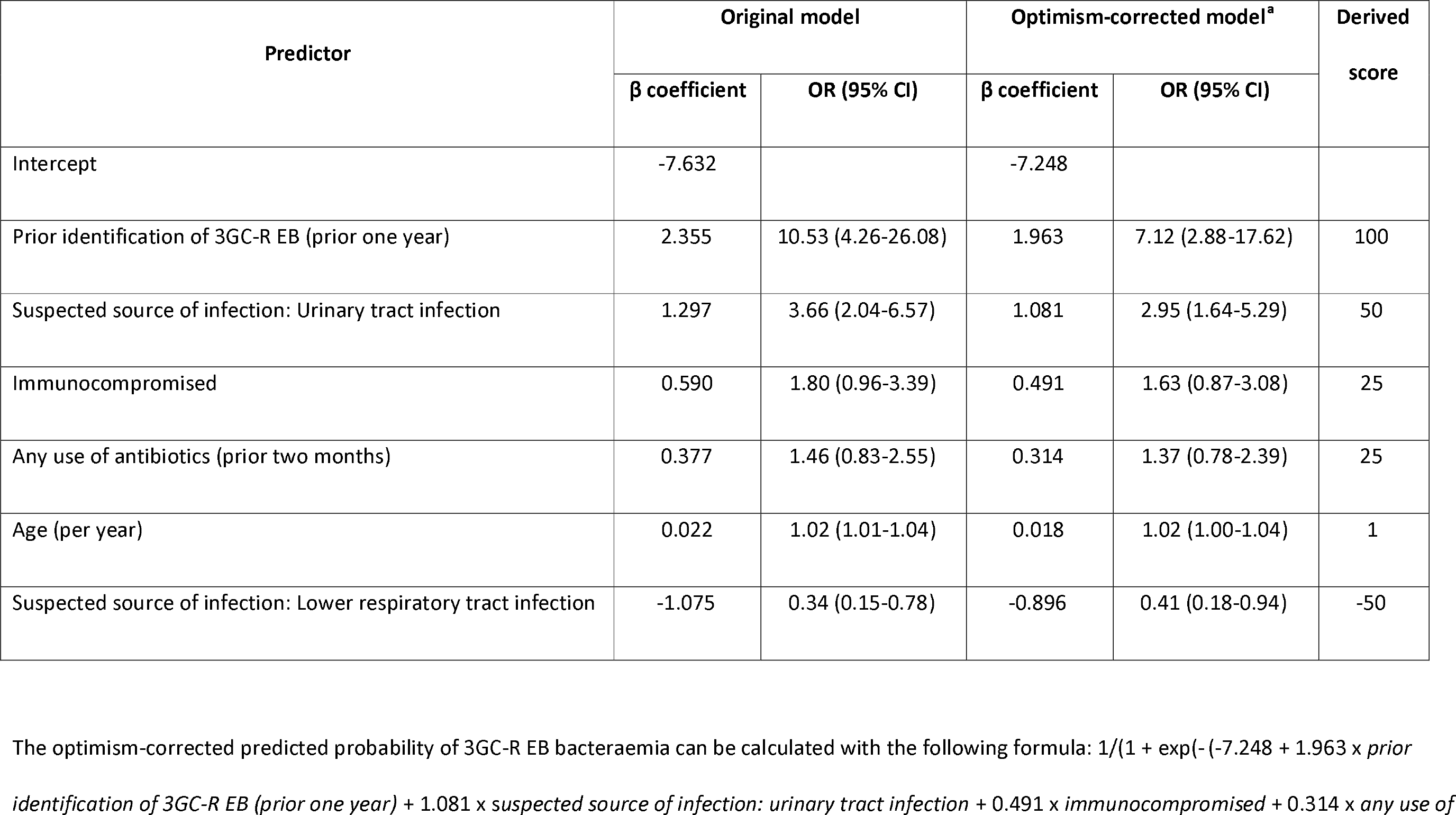
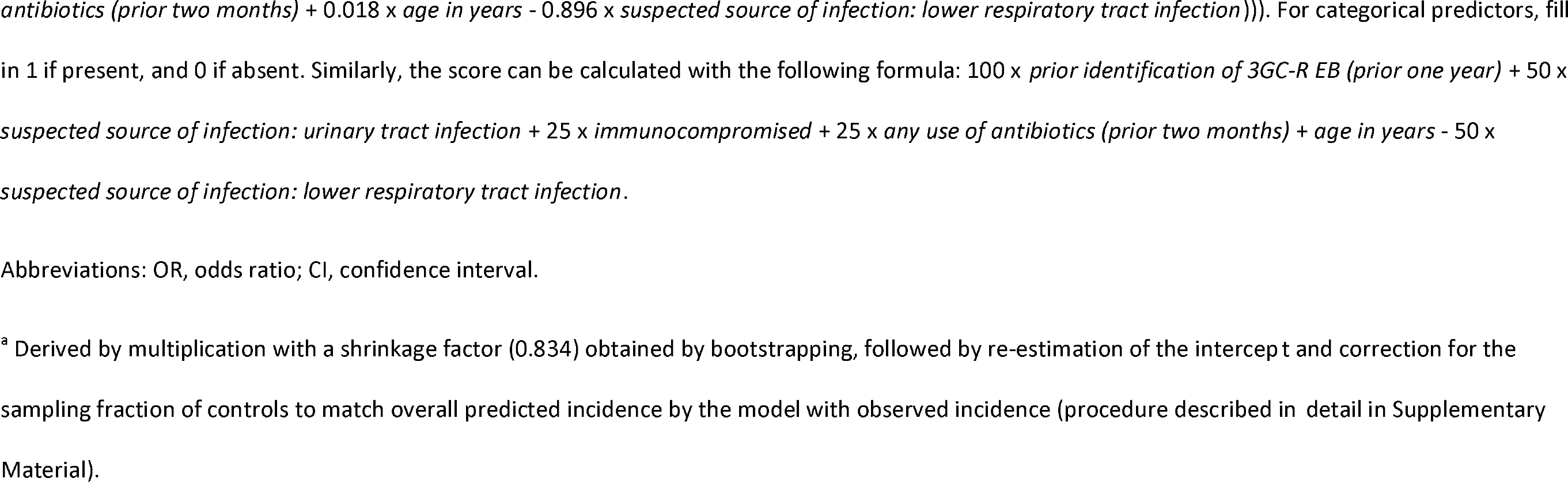
Regression model and scoring system for community-onset infection.

| Predictor | Original model |  | Optimism-corrected model<sup>a</sup> |  | Derived score |
| --- | --- | --- | --- | --- | --- |
|  | β coefficient | OR (95% CI) | β coefficient | OR (95% CI) |  |
| Intercept | -7.632 |  | -7.248 |  |  |
| Prior identification of 3GC-R EB (prior one year) | 2.355 | 10.53 (4.26-26.08) | 1.963 | 7.12 (2.88-17.62) | 100 |
| Suspected source of infection: Urinary tract infection | 1.297 | 3.66 (2.04-6.57) | 1.081 | 2.95 (1.64-5.29) | 50 |
| Immunocompromised | 0.590 | 1.80 (0.96-3.39) | 0.491 | 1.63 (0.87-3.08) | 25 |
| Any use of antibiotics (prior two months) | 0.377 | 1.46 (0.83-2.55) | 0.314 | 1.37 (0.78-2.39) | 25 |
| Age (per year) | 0.022 | 1.02 (1.01-1.04) | 0.018 | 1.02 (1.00-1.04) | 1 |
| Suspected source of infection: Lower respiratory tract infection | -1.075 | 0.34 (0.15-0.78) | -0.896 | 0.41 (0.18-0.94) | -50 |

Abbreviations: OR, odds ratio; CI, confidence interval.
<sup>a</sup> Derived by multiplication with a shrinkage factor (0.834) obtained by bootstrapping, followed by re-estimation of the intercept and correction for the sampling fraction of controls to match overall predicted incidence by the model with observed incidence (procedure described in detail in Supplementary Material).

The derived scoring system had a performance similar to the original model (Supplementary Figure 2a; C-statistic 0.807 (95% CI 0.756-0.855), not corrected for optimism). Table 3 and Figure 2a depict the trade-off between sensitivity and test positivity rate at different cutoffs for being at risk of 3GC-R EB bacteraemia. These can be contrasted to the fixed values for the *prior identification model* (sensitivity 24.4% and test positivity rate 2.8%), and the *two-predictor model* (sensitivity 53.9% and test positivity rate 21.5%).

**Figure 2.**
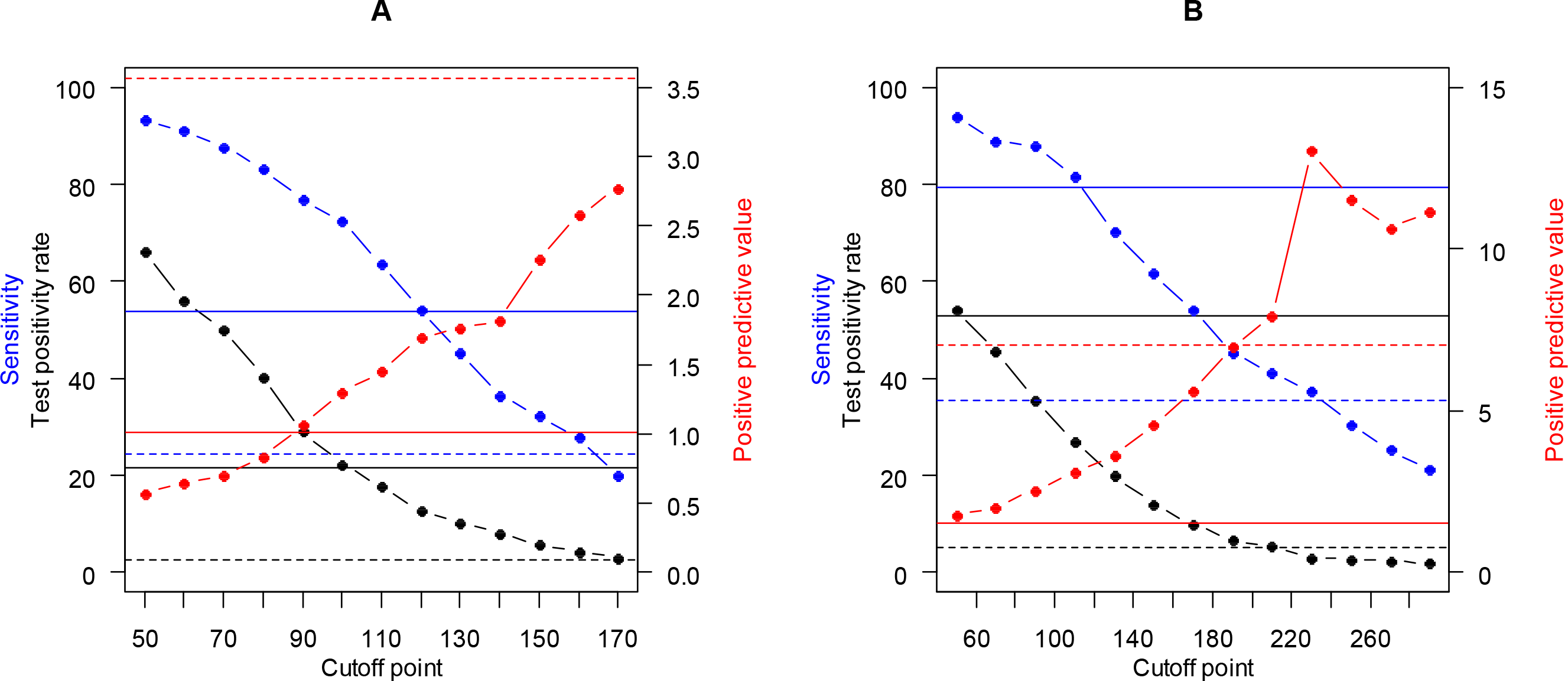
Performance of community-onset (A) and hospital-onset (B) scoring systems at different cutoff values. Figures show sensitivities (blue), test positivity rates (black), and positive predictive values (red) at different cutoffs for derived scoring systems from which patients are categorized as at risk of 3GC-R EB bacteraemia. These are compared to the (constant) sensitivities, test positivity rates, and positive predictive values for the basic two-predictor model (solid lines) and prior identification model (dashed lines). See Tables 3 and 6 for exact values at the score cutoffs.

**Table 3.** Performance of scoring system for community-onset infection.

|  | Score |  |  |  |  |  |  |  |  |  |  |  |  |  |  |
| --- | --- | --- | --- | --- | --- | --- | --- | --- | --- | --- | --- | --- | --- | --- | --- |
|  | -31<sup>a</sup> | 50 | 60 | 70 | 80 | 90 | 100 | 110 | 120 | 130 | 140 | 150 | 160 | 170 | 267<sup>b</sup> |
|  | Characteristics of interval [prior value, current value) |  |  |  |  |  |  |  |  |  |  |  |  |  |  |
| Proportion of population (%) |  | 33.9 | 10.1 | 6.0 | 9.7 | 11.3 | 6.7 | 4.7 | 4.8 | 2.5 | 2.2 | 2.3 | 1.4 | 1.4 | 2.9 |
| Probability of 3GC-R EB<br>bacteraemia (%) |  | 0.1 | 0.1 | 0.2 | 0.2 | 0.2 | 0.3 | 0.7 | 0.8 | 1.4 | 1.5 | 0.8 | 1.3 | 2.2 | 2.6 |
| | Characteristics of cutoff ≥ current value for classification as *at risk of 3GC-R EB bacteraemia* | | | | | | | | | | | | | | |
| Test positivity rate (%) |  | 66.1 | 56.0 | 50.0 | 40.3 | 29.0 | 22.4 | 17.7 | 12.8 | 10.3 | 8.1 | 5.7 | 4.3 | 2.9 | 0.0 |
| Sensitivity (%) |  | 93.2 | 91.0 | 87.8 | 83.3 | 76.8 | 72.3 | 63.7 | 54.3 | 45.2 | 36.6 | 32.2 | 27.8 | 20.0 | 1.1 |
| Specificity (%) |  | 34.0 | 44.1 | 50.1 | 59.9 | 71.2 | 77.8 | 82.5 | 87.3 | 89.8 | 92.1 | 94.4 | 95.8 | 97.2 | 100.0 |
| Positive predictive value (%) |  | 0.6 | 0.6 | 0.7 | 0.8 | 1.1 | 1.3 | 1.4 | 1.7 | 1.8 | 1.8 | 2.3 | 2.6 | 2.8 | 100.0 |
| Negative predictive value (%) |  | 99.9 | 99.9 | 99.9 | 99.9 | 99.9 | 99.9 | 99.8 | 99.8 | 99.8 | 99.7 | 99.7 | 99.7 | 99.7 | 99.6 |

<sup>a</sup> Minimum score within the study sample.
<sup>b</sup> Maximum score within the study sample.

For instance, patients with a score of 120 or higher would have a probability of 1.7% (positive predictive value) of having 3GC-R EB bacteraemia, and with this score as a cutoff 45.7% of all patients with 3GC-R EB bacteraemia would be missed (1 - sensitivity). This sensitivity (or proportion missed) is comparable to the simpler *two-predictor model*; however, the scoring system reduces eligibility for carbapenem use (test positivity rate) by 40%, from 21.5% to 12.8%.

Bootstrapping of the model indicated that when applying this cutoff in a future patient population some performance loss should be expected due to model optimism. The optimism-corrected sensitivity for future populations was 6.2 percentage points lower, whereas a change in prevalence was hardly noticeably (Table 4; please note that the percentages presented relate to the regression model, not to the score described in the paragraph above).

**Table 4.** Expected optimism when selecting probability cutoffs based on the sensitivity of the *two-predictor model*.

|  | Community-onset infection |  | Hospital-onset infection |  |
| --- | --- | --- | --- | --- |
|  | Two-predictor model | Probability cutoff 0.0067<sup>a</sup> | Two-predictor model | Probability cutoff 0.0086<sup>a</sup> |
|  | **Apparent performance in study sample** |  |  |  |
| **Sensitivity, % (95% CI)** | 53.9 (44.2-63.9) | 55.2 (43.7-63.7) | 79.3 (70.7-87.8) | 80.6 (71.8-88.8) |
| **Test positivity rate, % (95% CI)** | 21.5 (17.3-25.8) | 12.8 (9.8-16.7) | 52.8 (47.3-57.9) | 27.6 (22.6-32.1) |
|  | **Optimism-corrected performance<sup>b</sup>** |  |  |  |
| **Sensitivity, % (95% CI)** | 53.9<sup>c</sup> (44.2-63.9) | 49.0 (32.3-62.2) | 79.3<sup>c</sup> (70.7-87.8) | 75.3 (61.8-86.6) |
| **Test positivity rate, % (95% CI)** | 21.5<sup>c</sup> (17.3-25.8) | 13.2 (6.9-18.6) | 52.8<sup>c</sup> (47.3-57.9) | 28.2 (18.7-35.0) |

Abbreviations: CI, confidence interval.
<sup>a</sup> The predicted probability (as calculated by the regression model) from which patients are classified as *at risk of 3GC-R EB bacteraemia*. It is chosen such that the resulting sensitivity is as close as possible to the sensitivity of the *two-predictor model*.
<sup>b</sup> As obtained by bootstrapping described in Supplementary Material.
<sup>c</sup> Not affected by optimism due to pre-specification of models.

### Hospital-onset infection

The hospital-onset prediction model contained nine variables (Table 5), and also had adequate discrimination (C-statistic = 0.842 (95% CI 0.793-0.886), optimism-corrected 0.811 (95% CI 0.742-0.873) and calibration (Supplementary Figure 3).

**Table 5.** Regression model and scoring system for hospital-onset infection.

| Predictor | Original model |  | Optimism-corrected model<sup>a</sup> |  | Derived score |
| --- | --- | --- | --- | --- | --- |
| | $\beta$ coefficient | OR (95% CI) | $\beta$ coefficient | OR (95% CI) | |
| Intercept | -6.210 |  | -5.807 |  |  |
| Renal disease | 1.743 | 5.71 (2.24-14.55) | 1.372 | 3.94 (1.55-10.05) | 120 |
| Prior identification of 3GC-R EB (prior one year) | 1.718 | 5.57 (2.41-12.89) | 1.353 | 3.87 (1.67-8.95) | 120 |
| Any solid malignancy | 0.917 | 2.50 (1.29-4.87) | 0.722 | 2.06 (1.06-4.01) | 80 |
| Signs of hypoperfusion (at infection onset) | 0.646 | 1.91 (0.91-4.01) | 0.509 | 1.66 (0.79-3.49) | 40 |
| Surgical procedure (prior 30 days) | 0.564 | 1.76 (0.94-3.28) | 0.444 | 1.56 (0.84-2.91) | 40 |
| Central vascular catheter (at infection onset) | 0.533 | 1.70 (0.88-3.31) | 0.420 | 1.52 (0.78-2.95) | 40 |
| Use of cephalosporins (prior two months) | 0.527 | 1.69 (0.90-3.17) | 0.415 | 1.51 (0.81-2.83) | 40 |
| Length of hospital stay prior to infection (per day) | 0.014 | 1.01 (1.00-1.03) | 0.011 | 1.01 (1.00-1.03) | 1 |
| Suspected source of infection: Lower respiratory tract infection | -2.196 | 0.11 (0.04-0.35) | -1.729 | 0.18 (0.06-0.56) | -160 |

The optimism-corrected predicted probability of 3GC-R EB bacteraemia can be calculated with the following formula: $1/(1 + \exp(-(-5.807 + 1.372 \times \text{renal disease} + 1.353 \times \text{prior identification of 3GC-R EB (prior one year)} + 0.722 \times \text{any solid malignancy} + 0.509 \times \text{signs of hypoperfusion (at infection onset)} + 0.444 \times \text{surgical procedure (prior 30 days)} + 0.420 \times \text{central vascular catheter (at infection onset)} + 0.415 \times \text{use of cephalosporins (prior two months)} + 0.011 \times \text{length of hospital stay prior to infection in days} - 1.729 \times \text{suspected source of infection: lower respiratory tract infection})))$ . For categorical predictors, fill in 1 if present, and 0 if absent. Similarly, the score can be calculated with the following formula: $120 \times \text{renal disease} + 120 \times \text{prior identification of 3GC-R EB (prior one year)} + 80 \times \text{any solid malignancy} + 40 \times \text{signs of hypoperfusion (at infection onset)} + 40 \times \text{surgical procedure (prior 30 days)} + 40 \times \text{central vascular catheter (at infection onset)} + 40 \times \text{use of cephalosporins (prior two months)} + \text{length of hospital stay prior to infection in days} - 160 \times \text{suspected source of infection: lower respiratory tract infection}$ .
Abbreviations: OR, odds ratio; CI: confidence interval.
<sup>a</sup> Derived by multiplication with a shrinkage factor (0.788) obtained by bootstrapping, followed by re-estimation of the intercept and correction for the sampling fraction of controls to match overall predicted incidence by the model with observed incidence (procedure described in detail in Supplementary Material).

The derived scoring system again performed very similar to the original model (Supplementary Figure 2b; C-statistic 0.842 (95% CI 0.794-0.887), not corrected for optimism). In Table 6 and Figure 2b, sensitivity and test positivity rate at different scoring cutoffs are compared to the *prior identification model* (sensitivity 35.4% and test positivity rate 5.2%), and the *two-predictor model* (sensitivity 79.3% and test positivity rate 52.8%).

**Table 6.** Performance of scoring system for hospital-onset infection.

|  | Score |  |  |  |  |  |  |  |  |  |  |  |  |  |  |
| --- | --- | --- | --- | --- | --- | --- | --- | --- | --- | --- | --- | --- | --- | --- | --- |
|  | -159<sup>a</sup> | 50 | 70 | 90 | 110 | 130 | 150 | 170 | 190 | 210 | 230 | 250 | 270 | 290 | 432<sup>b</sup> |
|  | Characteristics of interval [prior value, current value) |  |  |  |  |  |  |  |  |  |  |  |  |  |  |
| Proportion of population (%) |  | 46.0 | 8.4 | 10.0 | 8.5 | 6.9 | 6.2 | 4.0 | 3.2 | 1.3 | 2.4 | 0.2 | 0.3 | 0.5 | 2.0 |
| Probability of 3GC-R EB<br>bacteraemia (%) |  | 0.1 | 0.6 | 0.1 | 0.8 | 1.7 | 1.4 | 2.0 | 2.8 | 3.1 | 1.6 | 30.2 | 19.3 | 8.7 | 10.6 |
| | Characteristics of cutoff ≥ current value for classification as *at risk of 3GC-R EB bacteraemia* | | | | | | | | | | | | | | |
| Test positivity rate (%) |  | 54.0 | 45.6 | 35.6 | 27.0 | 20.1 | 13.9 | 9.9 | 6.7 | 5.4 | 3.0 | 2.7 | 2.4 | 2.0 | 0.0 |
| Sensitivity (%) |  | 93.9 | 89.0 | 87.8 | 81.5 | 70.1 | 61.7 | 54.0 | 45.2 | 41.2 | 37.5 | 30.6 | 25.3 | 21.3 | 1.2 |
| Specificity (%) |  | 46.4 | 54.9 | 65.0 | 73.5 | 80.4 | 86.5 | 90.5 | 93.7 | 95.0 | 97.4 | 97.6 | 97.8 | 98.2 | 100.0 |
| Positive predictive value (%) |  | 1.8 | 2.0 | 2.5 | 3.1 | 3.6 | 4.6 | 5.6 | 7.0 | 7.9 | 13.0 | 11.5 | 10.6 | 11.1 | 100.0 |
| Negative predictive value (%) |  | 99.9 | 99.8 | 99.8 | 99.7 | 99.6 | 99.5 | 99.5 | 99.4 | 99.4 | 99.3 | 99.3 | 99.2 | 99.2 | 99.0 |

<sup>a</sup> Minimum score within the study sample.
<sup>b</sup> Maximum score within the study sample.

Patients with scores of 110 or higher have a 3.1% probability of 3GC-R EB bacteraemia, and with this cutoff 18.5% of all patients with 3GC-R EB bacteraemias would be missed, similarly to the *two-predictor model*. Yet, carbapenem eligibility would be reduced with 49% (27.0% vs. 52.8%). In this scenario, bootstrapping indicated that sensitivity in future patient populations should again be expected to be somewhat lower (-5.3%; Table 4).

## Discussion

We developed scoring systems to more accurately identify patients with bacteraemia caused by 3GC-R EB among those in whom empiric intravenous antibiotic therapy aimed at Gram-negatives is initiated. The scores consist of a limited number of clinical predictors that can easily be assessed based on the information available at the initial examination of a patient presenting with infection, before prescription of initial antibiotics, such as medical history, prior antibiotic usage, prior microbiology results, and infection characteristics. The calculated score can directly be converted to a probability that the patient suffers from 3GC-R EB bacteraemia, and depending on this probability, a decision can be made whether initial antibiotics should include coverage for 3GC-R EB or not. Implementing the scoring systems could improve appropriateness of empiric antibiotic therapy and reduce unnecessary use of broad-spectrum therapy. Compared to a basic model incorporating only prior 3GC-R EB identification and exposure to cephalosporins and/or fluoroquinolones, eligibility for empiric carbapenem use could be reduced by 40%-49% while maintaining a similar risk of missing patients with 3GC-R EB bacteraemia.

With a global emergence of antibiotic resistance, physicians must assess the risks of missing resistant causative pathogens when starting empiric antibiotic treatment [23]. Risk avoidance, albeit imaginable in many situations, is one of the driving forces for broad-spectrum antibiotic use, fuelling the global pandemic of antimicrobial resistance. Better prediction rules for infections caused by antibiotic-resistant pathogens are therefore needed. Prediction systems have been developed for Gram-negative bacteraemia in septic patients [24], carriage of or infection with ESBL-producing Enterobacteriaceae at hospital admission [1,25,26], and distinguishing bacteraemia with ESBL- or carbapenemase-producing pathogens from bacteraemia with susceptible Enterobacteriaceae [5–8].

Yet, guidance on incorporating the risk of 3GC-R EB in selecting empiric antibiotics is currently lacking. A recently published flow chart for initiating empiric therapy with a carbapenem in critically ill patients with suspected Gram-negative infection included predictors for 3GC-R EB carriage at hospital admission and in case of Enterobacteriaceae bacteraemia, without formal evaluation of performance [27]. For clarity, 3GC-R EB bacteraemia is a subset of Enterobacteriaceae bacteraemia, which is a subset of all bacteraemia episodes. Risk factors for any of these overarching categories may alter the probability of bacteraemia caused by antibiotic-resistant Enterobacteriaceae. This is corroborated by the strong predictive role of the suspected source of infection in our prediction models, which likely reflects the likelihood that Enterobacteriaceae play a role as causative pathogens. It emphasizes the need to select a clinically meaningful patient population when deriving a prediction rule. We therefore focused on all patients receiving their first dose of antibiotic therapy aimed at Enterobacteriaceae, rather than selecting patients that had, in retrospect, bacteraemia.

Due to the effect of including all patients with a clinical suspicion of infection, predicted probabilities of 3GC-R EB bacteraemia may seem low (0.4-1.0%). Yet, in a previous Dutch study, an 8.3% 3GC resistance rate among Enterobacteriaceae bacteraemia isolates resulted in a similarly low prior probability of 3GC-R EB bacteraemia in case of suspected Gram-negative infection (0.7%) [10]. Although our data originated from 2008-2010, the prevalence of 3GC resistance among Enterobacteriaceae only marginally increased in the Netherlands since then, and most Western European countries currently have similar prevalence rates of 3GC resistance among Enterobacteriace, namely between five and fifteen percent [28]. Model updating to reflect the local prevalence of resistance will generally improve calibration [22], but our model provides a useful universal backbone due to the incorporation of widely reported risk factors [29].

With the newly developed prediction rules, we aimed to achieve similar sensitivities as in existing prediction schemes, while at the same time reducing the proportion of patients eligible for broad-spectrum antibiotics (test-positives). This leads to diverging performance; for community-acquired infections we were “satisfied” with a sensitivity of 54.3%, where this figure was 81.5% for hospital-onset infections, and this yielded test-positive proportions of 12.8% and 27.0% for community-onset and hospital-onset infections, respectively. Yet, both prediction rules can also be used to increase sensitivity, which will – as a matter of fact – also increase the proportion of test-positivity. The optimal cut-off cannot be defined as each point has a different balance between the risk of overprescribing carbapenems and inappropriate empiric antibiotics.

That balance may be different for certain bug-drug combinations. For instance, the acceptance for a delay in adequate treatment of enterococcal bacteraemia may be different than for carbapenemase-producing Enterobacteriaceae, and might be different in a clinically stable than in a haemodynamically unstable patient [30]. Taking the long-term population effects of, for instance, carbapenem overuse into that equation is difficult, as these effects have not been sufficiently quantified [31], and depend on extraneous factors such as hospital hygiene and the baseline prevalence of carbapenem-resistant micro-organisms [32].

As expected, prior identification of 3GC-R EB was the strongest predictor in both models. In the Netherlands, screening for carriage is only practiced in intensive care units and for highly selected risk groups. Hence, identification was mostly based on previous clinical cultures, and an unknown proportion of actual 3GC-R EB carriers are classified as non-carriers. Naturally, more screening will further increase the sensitivity of this predictor for bacteraemia with 3GC-R EB. Yet, as infection rates among colonized patients are low [33,34], it is unsure whether positive predictive values of models will improve. In fact, if low-risk carriers would be identified by screening more frequently, positive predictive values might even decline.

Prior antibiotic use, on the other hand, had little predictive value in the community-onset model, and was not retained in the hospital-onset model. This seems to contradict the results from other studies. Yet, in such studies associations resulted from comparing infections with resistant Enterobacteriaceae to their sensitive counterparts [29], which exaggerates the role of antibiotic use [35].

We applied a nested case-control design for this study, implying that instead of analysing the full cohort, a representative subset of patients without 3GC-R EB (i.e. the control population) was analysed. The case population, however, (i.e. patients with 3GC-R EB) was analysed in full. This design was chosen for efficiency reasons, reducing the amount of data collection by 95% while accepting a small loss of precision. Knowing the size of the original cohort, we were able to extrapolate the case-control data to the full cohort, resulting in probabilities generalizable to clinical practice. Within the community-onset and hospital-onset cohorts, we matched on hospital to adjust for hospital-specific practices (independent of the incidence of 3GC-R EB) and on date to avoid effects of season-specific fluctuations in incidence and risk factors.

Our study has a limited sample size compared to the initial number of predictors studied. This may simultaneously lead to falsely rejecting predictive variables (a power problem) and selection of spurious predictors (overfitting resulting in overoptimism of model performance) [14]. We applied high p-value thresholds for variable retention in models to overcome our relatively low power, and internal validation by means of bootstrapping to quantify optimism in our model selection strategy. The latter resulted in optimism-adjusted odds ratios and C-statistics, giving insight in values expected when applying models to an external cohort. Expected performance loss when selecting specific probability cutoffs for clinical use has also been calculated (Table 4). Naturally, both models need prospective external validation before clinical implementation, for two reasons. First, even after shrinkage, optimism may still be present, as some steps could not be replicated in the bootstrap procedure, such as aggregation after observing similar associations with the outcome, simplification of continuous variables to linear predictors, and derivation of a scoring system. Second, the current study relied on data available in medical charts. We used pragmatic in- and exclusion criteria, which might not fully reflect intended clinical use, and as data collection was not blinded for outcome, information bias is not excluded. Moreover, potentially relevant predictors, especially for community-onset infection, such as international travel, animal contact, known colonization in household members, and dietary preferences could not be collected [29]. The same holds for determination of colonization pressure, which might be a relevant predictor for hospital-onset infections [32]. External validation studies are currently ongoing.

A study limitation is that the outcome was restricted to bacteraemic episodes, not including non-bacteraemic infections caused by 3GC-R EB, which are more common than bacteraemic infections in patients being empirically treated [10]. Yet, with an overall prevalence of <5% it is unlikely that these infections had a substantial impact on the composition of control groups. Future studies may consider classifying these infections as outcomes. However, due to the more benign course, initial treatment with carbapenems may not have a high priority in non-bacteraemic infections.

Another limitation of our study is that empiric coverage of 3GC-R EB is just one aspect of selection of appropriate empiric therapy. Other potential pathogens and resistance mechanisms, such as *Pseudomonas aeruginosa*, might justify alterations in empiric treatment even in the absence of risk factors for 3GC-R EB. In some countries, high incidences of infections with carbapenemase-producing Enterobacteriaceae may limit usefulness of our models. On the other hand, escape therapy for 3GC-R EB might not necessarily involve carbapenems, due to underlying resistance mechanisms other than ESBL, or favourable patterns of co-resistance. Ideally, frameworks for selecting empiric therapy should evaluate the probability of success of many different antibiotic agents. An example of such an approach is TREAT, an automated system for recommending antibiotic treatment based on, amongst others, patient and infection characteristics and local epidemiology [36]. TREAT can predict the presence of Gram-negative causative pathogens in infection with some accuracy [37], but performance with regard to resistant variants remains unknown. However, TREAT has not been widely adopted [38], and simple prediction rules may be easier to incorporate into clinical practice.

Furthermore, treating physicians incorporate more factors in their clinical decision making regarding empiric antibiotics than those provided by current risk stratification schemes in guidelines. In both this and our previous study [10], empiric carbapenem use was much lower than it would have been with full guideline adherence (Supplementary Table 4). As a result, achievable reductions in empiric carbapenem use in real life may be lower than anticipated in our study. Yet, we consider it important that antibiotic guidelines do not stimulate unnecessary broad-spectrum antibiotic use [39].

In conclusion, identification of patients with an infection caused by 3GC-R EB amongst all patients that need empiric antibiotic therapy remains a trade-off between acceptably low levels of unnecessary empiric carbapenem use and appropriate treatment in true 3GC-R EB bacteraemia cases. The prediction rules developed quantify this trade-off for patients that need empiric treatment, and might offer improvement in detecting such patients, compared to current international guidelines. As such, they provide useful starting points for optimizing empiric antibiotic strategies.

## Transparency declaration

W.C.R. has received research funding from Merck Sharp & Dohme (MSD). All other authors declare no conflicts of interest. This work was supported by a research grant from the Netherlands Organisation for Health Research and Development (project number 205200007) to H. S. M. A.

We would like to thank Evelien Stevenson and Tim Deelen for assistance with the data collection, and Maaike van Mourik and Loes de Bruin for assistance with the statistical analysis.

Results from this study were presented at the 26^th^ European Congress of Clinical Microbiology and Infectious Diseases, Amsterdam, the Netherlands (9-12 April 2016, E-poster eP0083), and at ASM Microbe 2016, Boston, MA, United States of America (16-20 June 2016, session 445). This manuscript is also available as an electronic preprint from www.bioRxiv.org (accessible via doi 10.1101/120550).

## References

[1] Tumbarello M, Trecarichi EM, Bassetti M, De Rosa FG, Spanu T, Di Meco E, et al. Identifying patients harboring extended-spectrum-beta-lactamase-producing Enterobacteriaceae on hospital admission: derivation and validation of a scoring system. Antimicrob Agents Chemother 2011;55:3485–90. doi:10.1128/AAC.00009-11.

[2] Platteel TN, Leverstein-van Hall M a., Cohen Stuart JW, Thijsen SFT, Mascini EM, van Hees BC, et al. Predicting carriage with extended-spectrum beta-lactamase-producing bacteria at hospital admission: a cross-sectional study. Clin Microbiol Infect 2015;21:141–6. doi:10.1016/j.cmi.2014.09.014.

[3] Pasricha J, Koessler T, Harbarth S, Schrenzel J, Camus V, Cohen G, et al. Carriage of extended-spectrum beta-lactamase-producing Enterobacteriacae among internal medicine patients in Switzerland. Antimicrob Resist Infect Control 2013;2:20. doi:10.1186/2047-2994-2-20.

[4] Shitrit P, Reisfeld S, Paitan Y, Gottesman B-S, Katzir M, Paul M, et al. Extended-spectrum beta-lactamase-producing Enterobacteriaceae carriage upon hospital admission: prevalence and risk factors. J Hosp Infect 2013;85:230–2. doi:10.1016/j.jhin.2013.07.014.

[5] Goodman KE, Lessler J, Cosgrove SE, Harris AD, Lautenbach E, Han JH, et al. A clinical decision tree to predict whether a bacteremic patient is infected with an extended-spectrum β-lactamase-producing organism. Clin Infect Dis 2016;63:896–903. doi:10.1093/cid/ciw425.

[6] Martin ET, Tansek R, Collins V, Hayakawa K, Abreu-Lanfranco O, Chopra T, et al. The carbapenem-resistant Enterobacteriaceae score: A bedside score to rule out infection with carbapenem-resistant Enterobacteriaceae among hospitalized patients. Am J Infect Control 2013;41:180–2. doi:10.1016/j.ajic.2012.02.036.

[7] Leibman V, Martin ET, Tal-Jasper R, Grin L, Hayakawa K, Shefler C, et al. Simple bedside score to optimize the time and the decision to initiate appropriate therapy for carbapenem-resistant Enterobacteriaceae. Ann Clin Microbiol Antimicrob 2015;14:1–5. doi:10.1186/s12941-015-0088-y.

[8] Augustine MR, Testerman TL, Justo JA, Bookstaver PB, Kohn J, Albrecht H, et al. Clinical Risk Score for Prediction of Extended-Spectrum β-Lactamase–Producing Enterobacteriaceae in Bloodstream Isolates. Infect Control Hosp Epidemiol 2016;38:1–7. doi:10.1017/ice.2016.292.

[9] Stichting Werkgroep Antibioticabeleid. SWAB guidelines for antibacterial therapy of adult patients with sepsis. Amsterdam, the Netherlands: 2010.

[10] Rottier WC, Bamberg YRP, Dorigo-Zetsma JW, van der Linden PD, Ammerlaan HSM, Bonten MJM. Predictive value of prior colonization and antibiotic use for third-generation cephalosporin-resistant Enterobacteriaceae bacteremia in patients with sepsis. Clin Infect Dis 2015;60:1622–30. doi:10.1093/cid/civ121.

[11] Pavlou M, Ambler G, Seaman SR, Guttmann O, Elliott P, King M, et al. How to develop a more accurate risk prediction model when there are few events. BMJ 2015;351:h3868. doi:10.1136/bmj.h3868.

[12] Rothman KJ, Greenland S, Lash TL. Case-Control Studies. In: Rothman KJ, Greenland S, Lash TL, editors. Modern Epidemiology. 3rd ed., Philadelphia, PA, USA: Lippincott Williams & Wilkins; 2008, p. 111–27.

[13] Collins GS, Reitsma JB, Altman DG, Moons KGM. Transparent Reporting of a multivariable prediction model for Individual Prognosis Or Diagnosis (TRIPOD): The TRIPOD Statement. Ann Intern Med 2015;162:55. doi:10.7326/M14-0697.

[14] Moons KGM, Altman DG, Reitsma JB, Ioannidis JP a., Macaskill P, Steyerberg EW, et al. Transparent Reporting of a multivariable prediction model for Individual Prognosis Or Diagnosis (TRIPOD): Explanation and Elaboration. Ann Intern Med 2015;162:W1. doi:10.7326/M14-0698.

[15] R Core Team. R: A Language and Environment for Statistical Computing. Vienna, Austria: 2015.

[16] Buuren S van, Groothuis-Oudshoorn K. mice: Multivariate Imputation by Chained Equations in R. J Stat Softw 2011;45. doi:10.18637/jss.v045.i03.

[17] Harrell Jr FE. rms: Regression Modeling Strategies 2016.

[18] Robin X, Turck N, Hainard A, Tiberti N, Lisacek F, Sanchez J-C, et al. pROC: an open-source package for R and S+ to analyze and compare ROC curves. BMC Bioinformatics 2011;12:77. doi:10.1186/1471-2105-12-77.

[19] Dahl DB. xtable: Export Tables to LaTeX or HTML 2016.

[20] White IR, Royston P, Wood AM. Multiple imputation using chained equations: Issues and guidance for practice. Stat Med 2011;30:377–99. doi:10.1002/sim.4067.

[21] Huang Y, Pepe MS. Assessing risk prediction models in case-control studies using semiparametric and nonparametric methods. Stat Med 2010;29:1391–410. doi:10.1002/sim.3876.

[22] Steyerberg EW. Clinical prediction models: A practical approach to development, validation, and updating. New York, NY: Springer Science+Business Media; 2009.

[23] Pogue JM, Kaye KS, Cohen DA, Marchaim D. Appropriate antimicrobial therapy in the era of multidrug-resistant human pathogens. Clin Microbiol Infect 2015;21:302–12. doi:10.1016/j.cmi.2014.12.025.

[24] Bates DW, Sands K, Miller E, Lanken PN, Hibberd PL, Graman PS, et al. Predicting bacteremia in patients with sepsis syndrome. J Infect Dis 1997;176:1538–51. doi:10.1086/514153.

[25] Slekovec C, Bertrand X, Leroy J, Faller J-P, Talon D, Hocquet D. Identifying patients harboring extended-spectrum-β-lactamase-producing Enterobacteriaceae on hospital admission is not that simple. Antimicrob Agents Chemother 2012;56:2218–9; author reply 2220. doi:10.1128/AAC.06376-ll.

[26] Johnson SW, Anderson DJ, May DB, Drew RH. Utility of a clinical risk factor scoring model in predicting infection with extended-spectrum β-lactamase-producing Enterobacteriaceae on hospital admission. Infect Control Hosp Epidemiol 2013;34:385–92. doi:10.1086/669858.

[27] Matteo B, Baño JR. Should we take into account ESBLs in empirical antibiotic treatment? Intensive Care Med 2016;42:2059–62. doi:10.1007/s00134-016-4599-6.

[28] European Centre for Disease Prevention and Control. Antimicrobial resistance surveillance in Europe 2015. Annual Report of the European Antimicrobial Resistance Surveillance Network (EARS-Net). Stockholm, Sweden: 2017.

[29] Trecarichi EM, Cauda R, Tumbarello M. Detecting risk and predicting patient mortality in patients with extended-spectrum β-lactamase-producing Enterobacteriaceae bloodstream infections. Future Microbiol 2012;7:1173–89. doi:10.2217/fmb.12.100.

[30] Paul M, Shani V, Muchtar E, Kariv G, Robenshtok E, Leibovici L. Systematic review and meta-analysis of the efficacy of appropriate empiric antibiotic therapy for sepsis. Antimicrob Agents Chemother 2010;54:4851–63. doi:10.1128/AAC.00627-10.

[31] Gharbi M, Moore LSP, Gilchrist M, Thomas CP, Bamford K, Brannigan ET, et al. Forecasting carbapenem resistance from antimicrobial consumption surveillance: Lessons learnt from an OXA-48-producing Klebsiella pneumoniae outbreak in a West London renal unit. Int J Antimicrob Agents 2015;46:150–6. doi:10.1016/j.ijantimicag.2015.03.005.

[32] Weinstein RA. Controlling antimicrobial resistance in hospitals: Infection control and use of antibiotics. Emerg Infect Dis 2001;7:188–92. doi:10.3201/eid0702.700188.

[33] Reddy P, Malczynski M, Obias A, Reiner S, Jin N, Huang J, et al. Reply to Blot et al. Clin Infect Dis 2008;46:482–482. doi:10.1086/526350.

[34] Ruppé E, Pitsch a, Tubach F, de Lastours V, Chau F, Pasquet B, et al. Clinical predictive values of extended-spectrum beta-lactamase carriage in patients admitted to medical wards. Eur J Clin Microbiol Infect Dis 2012;31:319–25. doi:10.1007/s10096-011-1313-z.

[35] Harris AD, Samore MH, Lipsitch M, Kaye KS, Perencevich E, Carmeli Y. Control-group selection importance in studies of antimicrobial resistance: examples applied to Pseudomonas aeruginosa, Enterococci, and Escherichia coli. Clin Infect Dis 2002;34:1558–63. doi:10.1086/340533.

[36] Leibovici L, Paul M, Nielsen AD, Tacconelli E, Andreassen S. The TREAT project: decision support and prediction using causal probabilistic networks. Int J Antimicrob Agents 2007;30 Suppl 1:S93–102. doi:10.1016/j.ijantimicag.2007.06.035.

[37] Paul M, Nielsen AD, Goldberg E, Andreassen S, Tacconelli E, Almanasreh N, et al. Prediction of specific pathogens in patients with sepsis: evaluation of TREAT, a computerized decision support system. J Antimicrob Chemother 2007;59:1204–7. doi:10.1093/jac/dkm107.

[38] Leibovici L. Decision support systems for antibiotic choices - better then their reputation? [presentation registered under #SY0303]. 27^th^ European Congress of Clinical Microbiology and Infectious Diseases, Vienna, Austria; 2017 April 23. Available from: https://www.escmid.org/escmid_publications/escmid_elibrary.

[39] Bonten MJM. Dangers and opportunities of guidelines in the data-free zone. Lancet Respir Med 2015;3:670–2. doi:10.1016/S2213-2600(15)00249-0.

[40] National Institute for Public Health and the Environment (RIVM). Beddencapaciteit ziekenhuizen 2008 n.d. http://www.zorgatlas.nl/zorg/ziekenhuiszorg/algemene-en-academische-ziekenhuizen/aanbod/beddencapaciteit-ziekenhuizen-2008/ (accessed March 17, 2016).

[41] Clinical and Laboratory Standards Institute. Performance standards for antimicrobial susceptibility testing; Seventeenth informational supplement. CLSI document M100-S17. Wayne, PA, USA: 2007.

[42] Clinical and Laboratory Standards Institute. Performance standards for antimicrobial susceptibility testing; Twenty-second informational supplement. CLSI document M100-S22. Wayne, PA, USA: 2012.

[43] ten Berg MJ, Huisman A, van den Bemt PMLA, Schobben AFAM, Egberts ACG, van Solinge WW. Linking laboratory and medication data: new opportunities for pharmacoepidemiological research. Clin Chem Lab Med 2007;45:13–9. doi:10.1515/CCLM.2007.009.

[44] Horan TC, Andrus M, Dudeck M a. CDC/NHSN surveillance definition of health care-associated infection and criteria for specific types of infections in the acute care setting. Am J Infect Control 2008;36:309–32. doi:10.1016/j.ajic.2008.03.002.

[45] Friedman ND, Kaye KS, Stout JE, McGarry SA, Trivette SL, Briggs JP, et al. Health care-associated bloodstream infections in adults: a reason to change the accepted definition of community-acquired infections. Ann Intern Med 2002;137:791–7.

[46] Charlson ME, Pompei P, Ales KL, MacKenzie CR. A new method of classifying prognostic comorbidity in longitudinal studies: development and validation. J Chronic Dis 1987;40:373–83.

[47] Levy MM, Fink MP, Marshall JC, Abraham E, Angus D, Cook D, et al. 2001 SCCM/ESICM/ACCP/ATS/SIS International Sepsis Definitions Conference. Intensive Care Med 2003;29:530–8. doi:10.1007/s00134-003-1662-x.

